# Highly efficient multiplex human T cell engineering without double-strand breaks using Cas9 base editors

**DOI:** 10.1101/482497

**Authors:** Beau R. Webber, Cara-lin Lonetree, Mitchell G. Kluesner, Matthew J. Johnson, Emily J. Pomeroy, Miechaleen D. Diers, Walker S. Lahr, Garrett Draper, Nicholas J. Slipek, Klaus N. Lovendahl, Amber McElroy, Wendy R. Gordon, Mark J. Osborn, Branden S. Moriarity

## Abstract

Chimeric antigen receptor engineered T cell (CAR-T) immunotherapy has shown efficacy against a subset of hematological malignancies^1,2^, yet its autologous nature and ineffectiveness against epithelial and solid cancers limit widespread application. To overcome these limitations, targeted nucleases have been used to disrupt checkpoint inhibitors and genes involved in alloreactivity^3–6^. However, the production of allogeneic, “off-the-shelf” T cells with enhanced function requires multiplex genome editing strategies that risk off-target effects, chromosomal rearrangements, and genotoxicity due to simultaneous double-strand break (DSB) induction at multiple loci^7–10^. Moreover, it has been well documented that DSBs are toxic lesions that can drive genetic instability^11,12^. Alternatively, CRISPR/Cas9 base editors afford programmable enzymatic nucleotide conversion at targeted loci without induction of DSBs^13,14^. We reasoned this technology could be used to knockout gene function in human T cells while minimizing safety concerns associated with current nuclease platforms. Through systematic reagent and dose optimization, we demonstrate highly efficient multiplex base editing and consequent protein knockout in primary human T cells at loci relevant to the generation of allogeneic CAR-T cells including the T cell receptor α constant (*TRAC*) locus, β-2 microglobulin (*B2M*), and programmed cell death 1 (*PDCD1*). Multiplex base edited T cells equipped with a CD19 CAR killed target cells more efficiently; and importantly, both DSB induction and translocation frequency were greatly reduced compared to cells engineered with Cas9 nuclease. Collectively, our results establish a novel multiplex gene editing platform to enhance both the safety and efficacy of engineered T cell-based immunotherapies.

---

Base editing has been previously used to induce premature stop (pmSTOP) codons for gene knockout in mice and in mammalian cells^15–18^. However, we reasoned that splice site disruption could have several advantages over induction of pmSTOP codons (**Supplemental Data 1**). For instance, stop codon readthrough has been shown to occur at frequencies up to 31% in some genes, and can be promoted under conditions of cellular stress^19,20^. Splice site editing mitigates this concern as it alters gene processing at the RNA level^21^, which is less likely to be bypassed at the translational level. Additionally, current base editors do not produce strict C to T edits, with even the most recent base editors producing up to 25% non-target editing (C to G/A)^22^. In the context of pmSTOP, non-target edits preclude premature stop codon formation, thereby lowering the efficiency of protein knockout, and instead create potentially undesirable amino acid changes.

To assess the performance of both pmSTOP introduction and splice-site disruption, we designed a panel of single guide RNAs (sgRNA) to convert amino acid codons to pmSTOPs or to disrupt splice donor (SD) and acceptor (SA) sequences within *PDCD1, TRAC*, and *B2M* (**Figure 1a, e, i; Supplemental Table 1**). Individual sgRNAs were co-delivered as chemically modified RNA oligonucleotides^23^ with first generation BE3^13^ or BE4^22^ mRNA to T cells by electroporation. Target C to T editing rates were assessed by Sanger sequencing and EditR, an analysis software developed by our group to expedite and economize analysis of base editing at the genetic level^24^ (baseeditr.com).

**Figure 1:**
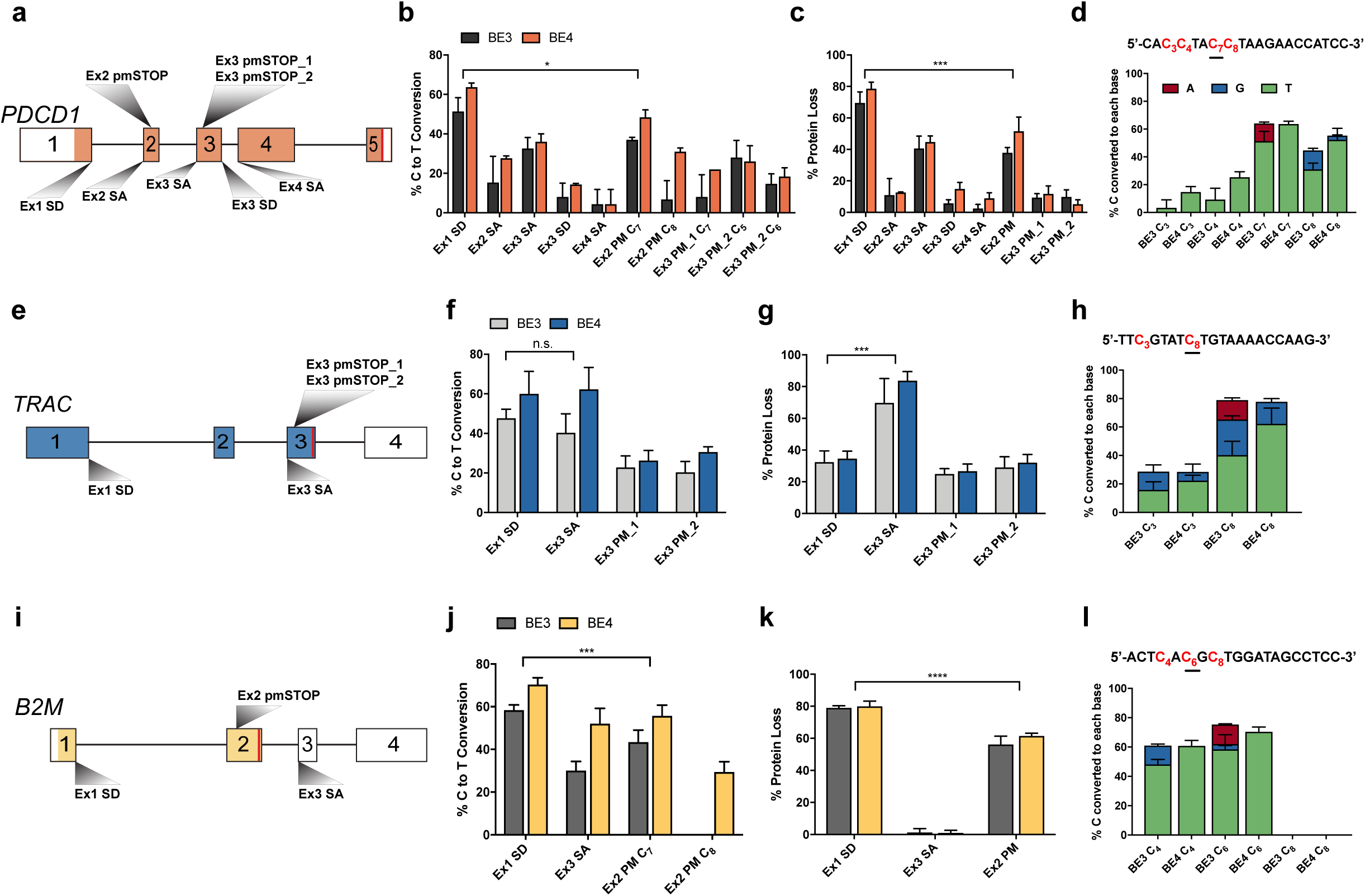
Assessment of guide RNA activity for gene disruption at *PDCD1, B2M*, and *TRAC*. Diagram of *PDCD1* locus indicating the relative locations of each sgRNA. (b) Quantification of C to T conversion of target base for each *PDCD1* sgRNA following co-delivery with either BE3 or BE4 mRNA as determined by EditR analysis of Sanger sequencing traces (*n=3* independent T cell donors). (c) PDCD1 protein knockout frequency after delivery of the indicated sgRNAs and either BE3 or BE4 mRNA as determined by flow cytometry (*n=3* independent T cell donors). (d) Quantification of C to T/A/G conversion at all Cs within the detected editing window (shown in red) of the *PDCD1* Ex1 SD sgRNA following co-delivery with either BE3 or BE4 mRNA as determined by EditR analysis of Sanger sequencing traces (*n=3* independent T cell donors). Underlined C indicates target nucleotide critical for proper splicing. (e) Diagram of *TRAC* locus indicating the relative locations of each sgRNA. (f) Quantification of C to T conversion at target base for each *TRAC* sgRNA following co-delivery with either BE3 or BE4 mRNA as determined by EditR analysis of Sanger sequencing traces (*n=3* independent T cell donors). (g) TRAC protein knockout frequency after delivery of the indicated sgRNAs and either BE3 or BE4 mRNA as determined by flow cytometry for CD3 loss (*n=3* independent T cell donors). (h) Quantification of C to T/A/G conversion at all cytosines within the detected editing window (shown in red) of the *TRAC* Ex3 SA sgRNA following co-delivery with either BE3 or BE4 mRNA as determined by EditR analysis of Sanger sequencing traces (*n=3* independent T cell donors). (i) Diagram of *B2M* locus indicating the relative locations of each sgRNA. (j) Quantification of C to T conversion of target base for each *B2M* sgRNA following co-delivery of either BE3 or BE4 mRNA as determined by EditR analysis of Sanger sequencing traces (*n=3 independent T cell donors*). (k) B2M protein knockout frequency after delivery of the indicated sgRNAs and either BE3 or BE4 mRNA as determined by flow cytometry for B2M loss (*n=3* independent T cell donors). (l) Quantification of C to T/A/G conversion at all cytosines within the detected editing window (shown in red) of the *B2M* Ex1 SD sgRNA following co-delivery with either BE3 or BE4 mRNA as determined by EditR analysis of Sanger sequencing traces (*data represented as mean ± SD, n=3 independent biological T cell donors*). *P*-values calculated by Student’s paired two-tailed t-test between the highest-editing guide and the second highest-editing treatment (n.s. *P* > 0.05, * *P* ≤ 0.05, ** *P* ≤ 0.01, *** *P* ≤ 0.001, **** *P* ≤ 0.0001).

First, we targeted the checkpoint gene *PDCD1* (PD-1) by designing eight sgRNAs; three of which were predicted to introduce pmSTOP codons, two targeted disruption of SD sites (GT:CA), and three targeted disruption of SA sites (AG:TC) (**Figure 1a**). We found that co-delivery of sgRNAs with BE3 or BE4 mRNA mediated measurable editing of target Cs at all target loci, with several candidate sgRNAs exhibiting significantly higher rates of editing than others (**Figure 1b, Supplemental Data 2**). Specifically, we found that targeting the SD site of *PDCD1* exon 1 resulted in the highest rate of target C to T editing with both BE3 (51.3 ± 7.0%, M ± SD) and BE4 (63.7 ± 2.1%) mRNA (**Figure 1b**). The next two most efficient sgRNAs targeted the exon 3 SA site (32.6 ± 5.5% for BE3; 36.0 ± 4.0% for BE4) and a candidate pmSTOP site in exon 2 (37.1 ± 1.2% for BE3; 48.5 ± 3.7% for BE4) (**Figure 1b**). To determine whether genetic editing results in protein loss we assessed expression of PD-1 protein by flow cytometry. Concordant with our genetic analysis, targeting *PDCD1* exon 1 SD resulted in the highest rate of protein loss (69.5 ± 7.0% for BE3; 78.6 ± 4.1% for BE4), followed by exon 3 SA (40.6 ± 7.8% for BE3; 44.7 ± 3.8% for BE4), and exon 2 pmSTOP (37.9 ± 3.4% for BE3; 51.5 ± 9.0% for BE4) (**Figure 1c**).

Informed by our *PDCD1* results, we designed a focused panel of sgRNAs targeting *TRAC* (**Figure 1e**). Here we found that C to T conversion was highest at the exon 1 SD site (47.6 ± 4.6% for BE3; 60.0 ± 11.3% for BE4) and exon 3 SA site (40.3 ± 9.7% for BE3; 62.3 ± 11.0% for BE4), with BE4 exhibiting higher editing rates than BE3 at each target (**Figure 1f**). Efficient editing was also observed at two pmSTOP candidate sites in exon 3, albeit at lower efficiencies than that of either splice-site disrupting sgRNA (**Figure 1f**). Both the exon 1 SD and exon 3 SA sites were edited at similar frequencies, yet disruption of the exon 3 SA site resulted in the highest rate of TCR disruption as measured by loss of cell-surface CD3 expression (69 ± 15.3% for BE3; 83.7 ± 5.8% for BE4) (**Figure 1g**).

We next targeted *B2M* using a similar strategy (**Figure 1i**). BE4 mRNA delivered with an sgRNA targeting the exon 1 SD site showed the most efficient C to T conversion of the target base (58.3 ± 2.5% for BE3; 70.3 ± 3.2% for BE4) (**Figure 1j**), resulting in efficient knockout of B2M protein (79.1 ± 1.3% for BE3; 80.0 ± 3.2% for BE4) (**Figure 1k**). We also identified a candidate pmSTOP site in exon 2 that resulted in relatively efficient C to T editing (43.3 ± 5.7% for BE3; 55.7 ± 5.0% for BE4), and protein knockout (56.2 ± 5.1% for BE3; 61.5 ± 1.8% for BE4) (**Figure 1j, k**). Notably, targeting the SA site of noncoding exon 3 produced efficient C to T editing but did not result in a detectable reduction in protein expression (**Figure 1j, k**).

Non-target editing (i.e. C to A or G) has been reported for BE3^13^ and is reduced with BE4, which contains a second uracil glycosylase inhibitor (UGI) fused in series at the C-terminus^22^. We evaluated non-target editing rates for all Cs within the editing window (predominantly bases 4-8 of protospacer) of our most efficient sgRNAs with BE3 and BE4. As expected, BE4 showed reduced non-target editing compared to BE3 at all loci (−14% ± 6.6%, *P* < 2.2e-16, Paired one-way t-test) (**Figure 1d, h, l**; **Supplemental Data 2**). Despite having only nickase function, low-level indel formation has been observed with both BE3 and BE4^13,22^. Thus, we used next-generation sequencing (NGS) to measure indel frequency at all target sites after editing (**Supplemental Data 3**). Indels were detectable with both BE3 and BE4 at levels that varied based on target site. Consistent with prior publications, BE4 exhibited an overall reduced indel frequency (−4.8% ± 6.1%, *P* < 4.6e-16, Paired one-way t-test) (**Supplemental Data 3**)^22^.

Toward our goal of validating a multiplex editing strategy that could be utilized to generate allogeneic, “off-the-shelf” T cells with enhanced function, we co-delivered our top sgRNA for each gene along with first-generation BE3 or BE4 mRNA. Surprisingly, editing efficiency at each target was substantially reduced for both BE3 and BE4 when delivered in a multiplex setting (**Supplemental Data 4**). To determine if the reduced editing efficiency was due to low protein levels, we delivered equal doses of BE3, BE4, and nuclease active *Streptococcus pyogenes* Cas9 (SpCas9) mRNA to T cells and measured protein expression at 24hrs and 48hrs after electroporation. Strikingly, while Cas9 protein expression was readily detectable at these time points, BE3 and BE4 protein were undetectable (**Supplemental Data 5**). To address this issue, we first delivered BE3 and BE4 mRNA at a dose 2x higher (3 µg) than that used in our initial multiplex experiments (1.5 µg). This strategy improved editing efficiency at each locus, but the efficiencies were still lower than those observed in our single gene targeting experiments (**Figure 2a**).

**Figure 2:**
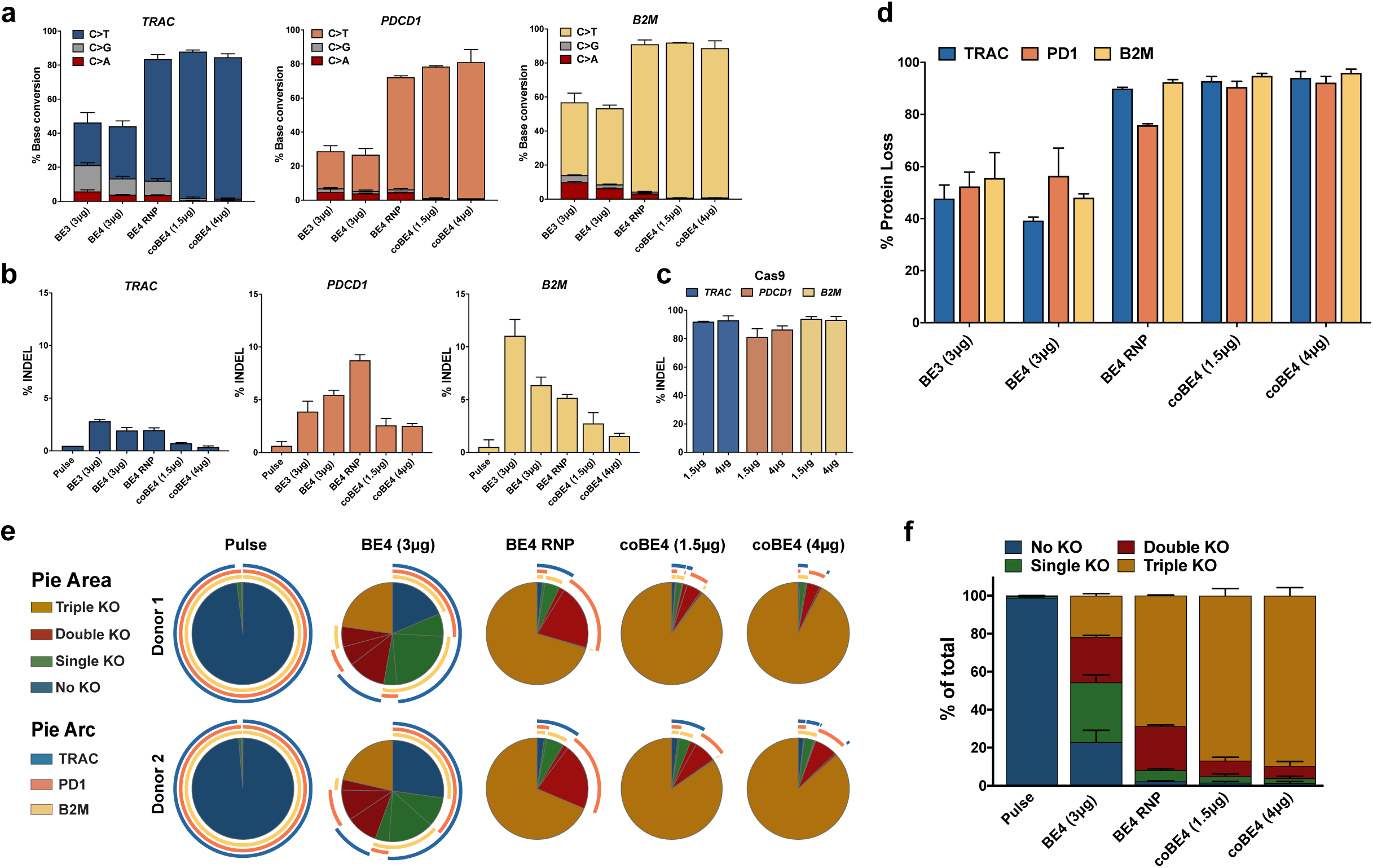
Optimization of multiplex editing using optimal sgRNAs (TRAC Ex3 SA, B2M Ex1 SD, and PD-1 Ex1 SD). (a) Conversion frequency of target cytosine to all other bases at *TRAC, PDCD1*, and *B2M* as analyzed by NGS following co-delivery of three target sgRNA with first-generation BE3 (BE3) or BE4 (BE4) mRNA delivered at 3 µg dose; BE4 protein complexed with sgRNA (BE4 RNP); or codon optimized BE4 mRNA (coBE4) delivered at 1.5 µg or 4 µg doses. Indel frequency at *TRAC, PDCD1*, and *B2M* as analyzed by NGS following delivery of three target sgRNA and first-generation BE3 (BE3) or BE4 (BE4) mRNA delivered at 3 µg dose; BE4 protein complexed with sgRNA (BE4 RNP); or codon optimized BE4 mRNA (coBE4) delivered at 1.5 µg or 4 µg doses. (c) Indel frequency at *TRAC, PDCD1*, and *B2M* as analyzed by NGS following co-delivery of three target sgRNA and SpCas9 nuclease mRNA at 1.5 µg or 4 µg dose. (d) Frequency of TRAC, PDCD1, and B2M protein loss measured by flow cytometry seven days after delivery of three target sgRNA and first-generation BE3 (BE3) and BE4 (BE4) mRNA delivered at 3 µg dose; BE4 protein complexed with sgRNA (BE4 RNP); and codon optimized BE4 mRNA (coBE4) delivered at 1.5 µg dose and 4 µg dose. (e) SPICE representation of multiplex flow cytometric analysis performed seven days post electroporation. (f) Quantification of fractions of WT, single, double, and triple gene KO. Data represented as mean ± SD, *n*=2 two independent biological T cell donors.

During the course of these experiments, independent reports emerged identifying problems related to the use of first-generation BE3 and BE4 expression vectors that severely reduce both transcriptional and translational efficiency in human cells^17,25^. To circumvent these issues, we delivered purified BE4 protein as a ribonucleoprotein (RNP) complex with our most effective sgRNA for each target. By optimizing our electroporation protocol for RNP delivery, we found that BE4 RNP mediated improved editing efficiency over a 2x dose of first-generation BE4 mRNA (**Figure 2c**). Next, we codon optimized the sequence of BE4 (coBE4) and delivered mRNA at both our standard dose (1.5 µg) and a higher dose (4 µg) with all three of our optimal sgRNAs. At both mRNA doses, we achieved substantially higher rates of multiplex target C to T editing at all three loci across multiple independent T cell donors, exceeding 90% in some instances (**Figure 2a**). Non-target editing observed with first-generation BE3 and BE4 mRNA was reduced slightly when using BE4 RNP, but even further reduced with both doses of coBE4 mRNA (**Figure 2a**). We next evaluated the rate of indel formation at each target site after multiplex base editing and, in accordance with previous studies, found lower rates of indel formation at each site with all forms of BE4 compared to BE3 and Cas9 nuclease **(Figure 2b, c)**. Both low and high doses of coBE4 mRNA exhibited the lowest overall frequency of indel formation at all sites examined **(Figure 2b)**. Multiplex protein knockout was analyzed for each target gene by flow cytometry, and the frequency of protein loss closely correlated with genetic editing frequencies **(Figure 2d)**. BE4 RNP demonstrated more efficient protein knockout than first-generation BE3 and BE4 mRNA, yet coBE4 mRNA was most efficient, exceeding 90% protein loss for each gene at both low and high mRNA doses **(Figure 2d, Supplemental Data 6).** A key consideration of multiplex editing is the resultant proportion of cells carrying each potential combination of gene knockout. To better understand this phenomenon in our experiments, we evaluated protein expression of all target genes simultaneously by flow cytometry and used SPICE analysis to determine the proportion of individual cells having no knockout, single gene knockout, double gene knockout, or triple gene knockout; as well as the combination of proteins lost within each of these fractions (**Figure 2e**). While first-generation BE4 mRNA generated an endpoint cell population with a diverse combination of knockout phenotypes, the frequency of triple knockout cells was low (21.9 ± 1.1%). The proportion of triple knockout cells was substantially higher using BE4 RNP (68.6 ± 0.37%), and even further increased with coBE4 mRNA at 1.5 µg (86.6 ± 3.75%) and 4 µg (89.57 ± 4.2%) (**Figure 2f**).

Nuclease-mediated multiplex editing has been reported to generate undesired translocations in human T cells^3^. As base editing substantially reduces the frequency of DSB formation, we reasoned that translocations should likewise be reduced using our base editing approach. To test our hypothesis, we used droplet digital PCR (ddPCR) assays that span the junction of several possible translocation outcomes **(Figure 3a).** Following co-delivery of our three optimal sgRNA with either spCas9 nuclease RNP or mRNA, we were able to detect three translocation outcomes: *B2M*:*TRAC*, *PDCD1*:*B2M*, and *PDCD1*:*TRAC* **(Figure 3b, c, d)**. In all cases, spCas9 mRNA resulted in the highest rate of translocation frequency, with translocations between *B2M* and *TRAC* being most frequent (1.57 ± 0.07%) **(Figure 3b, c, d)**. In stark contrast, the *B2M*:*TRAC* and *PDCD1*:*TRAC* outcomes were not detected in cell populations receiving BE4 RNP or either dose of coBE4 mRNA with our optimal sgRNAs **(Figure 3b, c, d)**. In a single replicate from one donor, the *PDCD1*:*B2M* assay gave rise to two positive droplets with low-dose coBE4 mRNA (calculated frequency = 0.003 ± 0.006%). Because no positive droplets were detected with BE4 RNP or high-dose coBE4 mRNA, these may be artifactual (**Figure 3c).**

**Figure 3:**
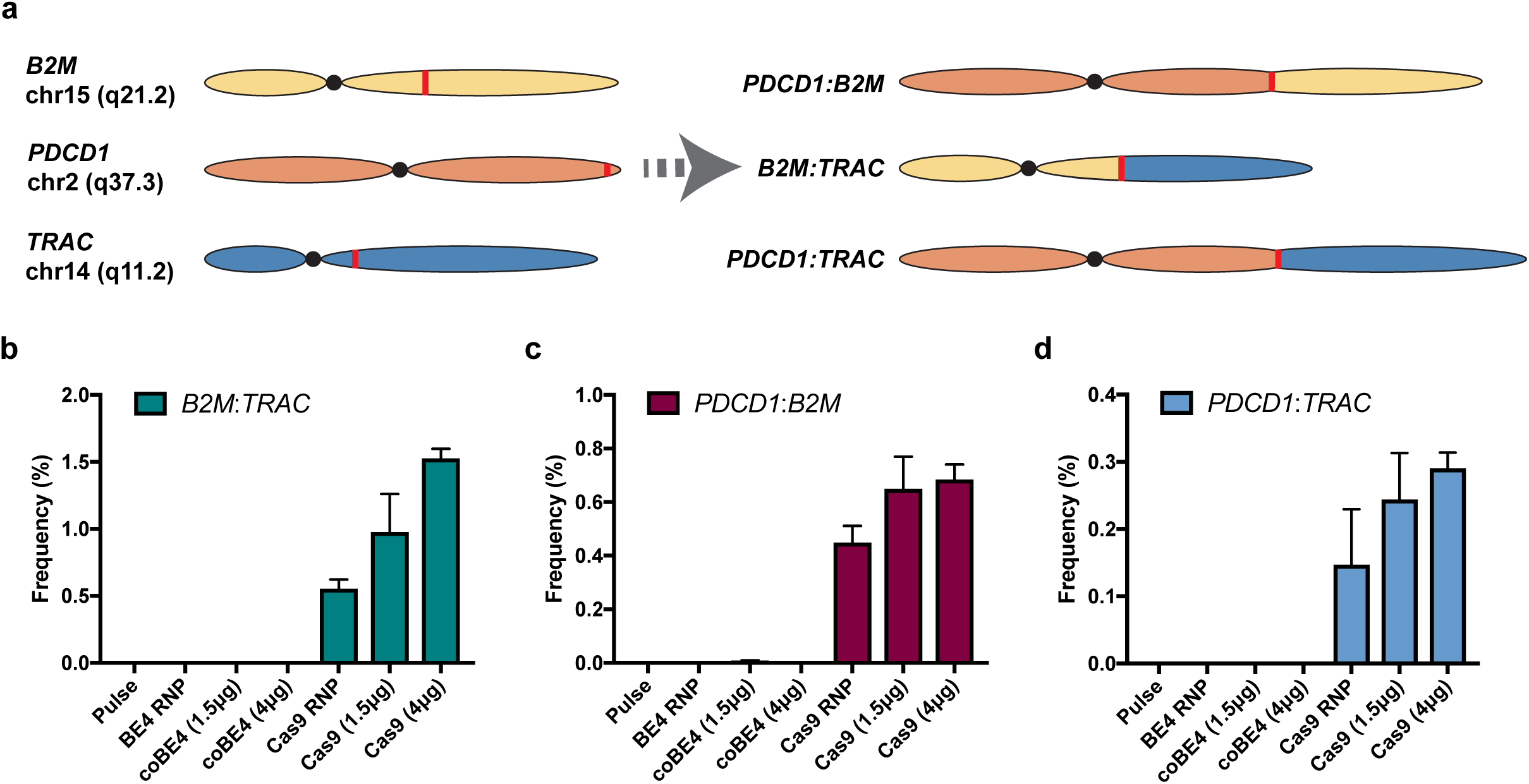
Translocation frequencies in multiplex edited T cells. (a) Diagram of assayed translocation outcomes. (b) Quantification of *B2M:TRAC* translocation outcome by ddPCR. (c) Quantification of *PDCD1:B2M* translocation outcome by ddPCR. (d) Quantification of *PDCD1:TRAC* translocation outcome by ddPCR. Data represented as mean ± SD. Assays run in technical duplicate across *n=2* independent biological T cell donors.

Off-target (OT) DSB induction is another important challenge facing nuclease platforms^26^. To determine the specificity of our optimal sgRNAs, we delivered each individually with spCas9 nuclease or BE4 mRNA and evaluated editing at the top 10 predicted OT sites by NGS (**Supplemental Table 2**). No editing was observed at any of the predicted *B2M* or *TRAC* OT sites in either the spCas9 nuclease or BE4 treatment conditions (**Supplemental Data 7**). At the predicted *PDCD1* OT sites we observed a single OT edit with an indel frequency of 13.0% using spCas9 mRNA (**Supplemental Data 7**). Strikingly, C to T editing at this site was only 0.9% with BE4 mRNA, and indel formation was near the low detection limit of our assay (0.2%) (**Supplemental Data 7**).

We next sought to determine whether multiplex knockout T cells generated using our base editing strategy retain cytokine functionality and are capable of mediating target cell killing when equipped with a CAR. We performed phenotypic evaluation of both electroporation pulse control and coBE4 knockout T cells with and without a CD19-specific CAR by analyzing markers of differentiation^27^. Both untransduced and CAR-transduced T cells exhibited similar differentiation phenotypes, with the fractions of effector and memory populations similar between control and coBE4 knockout T cells (**Figure 4a**). CAR transduction and cell expansion were also comparable between pulse and coBE4 mRNA groups (**Supplemental Data 8**). Following activation, a high frequency of both untransduced and CAR-transduced coBE4 knockout T cells exhibited robust production of cytokines IL-2, TNFα, and IFNγ (**Figure 4b**). Cytokine polyfunctionality was similarly retained following the multiplex editing process (**Figure 4c**). Collectively, these data demonstrate that multiplex coBE4 editing combined with CAR transduction did not negatively impact T cell phenotype or function. Finally, to determine if coBE4 knockout T cells equipped with the CD19 CAR retained the ability to kill target cells, we conducted *in vitro* co-culture assays with non-target CD19^neg^/PD-L1^neg^ K562; target CD19^pos^/PD-L1^neg^ Raji; and target CD19^pos^/PD-L1^pos^ Raji engineered to overexpress PD-L1, which would normally act to inhibit killing by T cells expressing cell surface PD-1. Both control and coBE4 knockout T cells mediated specific killing of CD19^pos^ but not CD19^neg^ target cells (**Figure 4d**). However, only coBE4 knockout T cells were able to achieve significant killing of CD19^pos^/PD-L1^pos^ target cells, with the efficiency of killing equivalent to that of CD19^pos^/PD-L1^neg^ target cells (**Figure 4d**).

**Figure 4:**
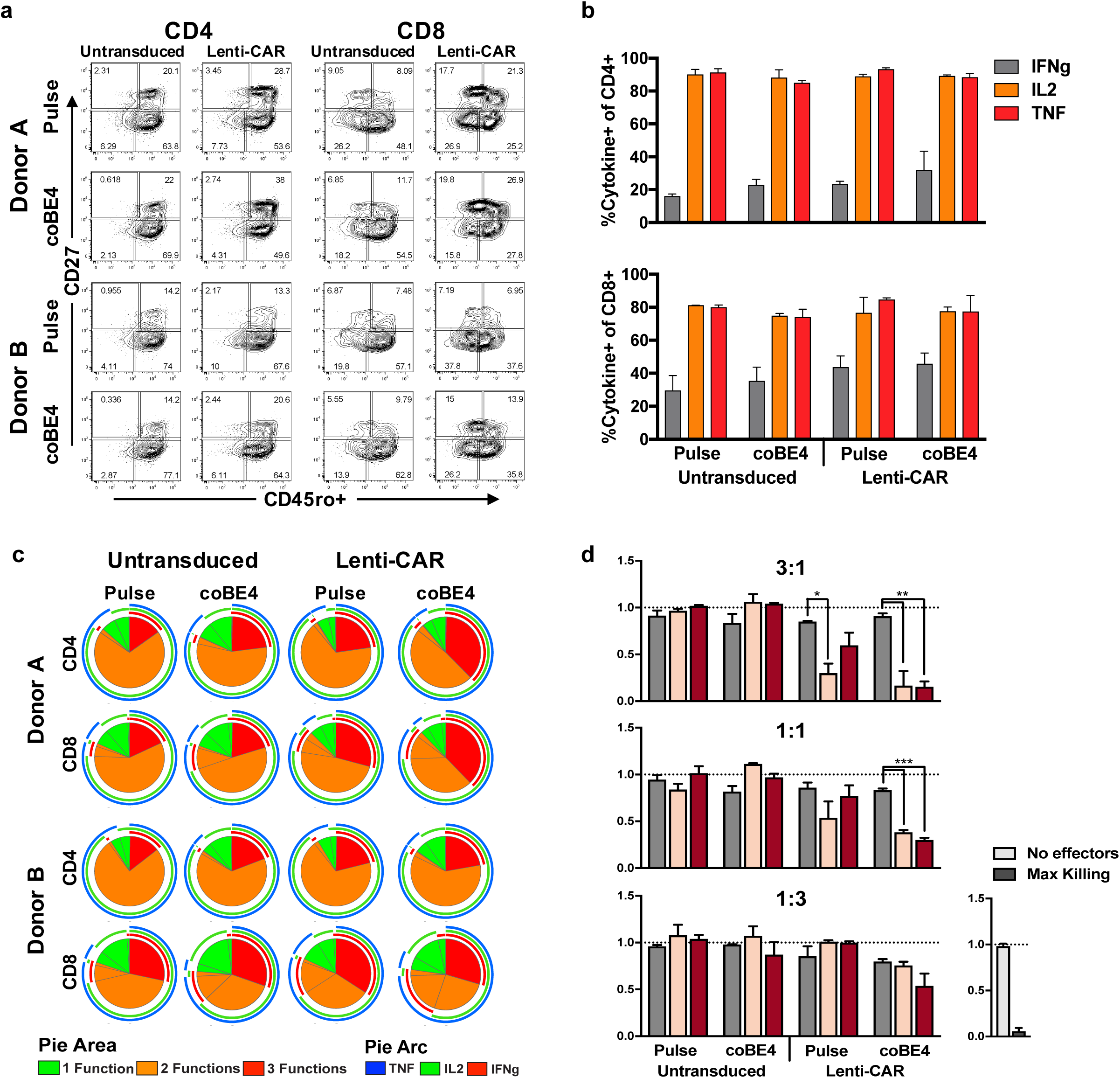
Function of multiplex edited T cells. (a) Expression of the memory marker CD27 and CD45ro following editing and expansion. Production of cytokines individually (b) and in combination (c) by CD4 and CD8 T cells following activation. (d) Ability of T cells to kill CD19^neg^ K562, CD19^pos^ Raji cells, or CD19^pos/^PD-L1^pos^ Raji cells as measured by luciferase luminescence assay following coculture with T cells. Graph titles indicate E:T ratio. Data represented as mean ± SD, with assays run in triplicate in two independent biological T cell donors. (n.s. *P* > 0.05, * *P* ≤ 0.05, ** *P* ≤ 0.01, *** *P* ≤ 0.001, **** *P* ≤ 0.0001).

As we come to better understand the requirements for successful cell-based immunotherapy and gene therapy, and as enthusiasm grows for the production of universal, allogeneic cells, highly multiplexed gene editing will likely become more commonplace. However, it has been well documented that DSBs are toxic lesions that can drive genomic instability and cell death^11,12^. This is a lesser concern when engineering cells for research but could lead to transformation or reduced function when gene editing cells for therapeutic use. Our concerns surrounding DSBs are further heightened in the context of multiplex gene editing where multiple, simultaneous DSBs can compound toxicity and increase the potential for detrimental translocations. To overcome these issues, we have implemented the use of base editor technology for multiplex T cell engineering and demonstrate that splice site disruption through base editing offers an efficient and safer approach compared to the use of DSB-inducing targeted nucleases.

Interestingly, we find both higher rates of non-target editing and indel formation when using BE4 RNP compared to coBE4 expressed from transfected mRNA. This observation may be due to the extended BE4 residence time achieved when expressed at high levels from a stable mRNA as opposed to direct BE4 protein delivery. As even free UGI has been shown to reduce both indel frequency and non-target editing in the context of BE3^28^, the extended residence time achieved by mRNA delivery may allow BE4 UGI domains additional capacity to mitigate DSB formation and non-target editing^28^.

In our current study we utilized lentiviral delivery of CD19-specific CAR, which is the current industry standard in CAR-T therapy. However, this approach has many drawbacks, including the risk of insertional mutagenesis, variable CAR expression, and gene silencing^29–31^. To overcome these issues, a number of groups have demonstrated high efficiency, site-specific integration using Cas9 nuclease along with rAAV-delivered DNA donor templates for homologous recombination (HR). This raises the possibility that BE4 could be deployed to safely and efficiently knockout multiple genes with simultaneous introduction of therapeutic transgenes in a site-specific fashion using rAAV and Cas9 orthologs, such as *Staphylococcus aureus* Cas9 (SaCas9) or *Francisella novella* Cas9 (FnCas9)^32,33^. The application of Cas9 orthologs would allow for simultaneous use of distinct sgRNAs specific to BE4 and Cas9 nuclease without concerns of cross-utilization. Alternatively, it has been demonstrated that a DNA nick can be used to stimulate HR using naked DNA or rAAV as a DNA donor molecule, albeit with lower efficiency^34^. This provocatively highlights the potential of BE4 to mediate gene knockout through deaminase activity, while simultaneously mediating HR through its nickase function. In this scenario, the sgRNA binding sites may require an absence of cytosines within the base editing window to prevent loss of Cas9 binding due to sequence changes through C to T conversion.

One notable difference between the use of base editors and targeted nucleases is the number of potential outcomes from the editing event. Nuclease-mediated DSBs are repaired through the highly variable non-homologous end joining (NHEJ) pathway, resulting in a spectrum of indels; some of which will not introduce frame-shift mutations and will thus have unknown significance to gene expression and function. Alternatively, our base editing approach has a limited number of outcomes, all resulting in the loss of function of the native splice donor or acceptor, even when considering non-target editing. Yet it is important to consider that disruption of the native splice site may not always result in a nonfunctional product, given that alternative or cryptic splicing could maintain the biological function of a gene.

Translocation analysis using small ddPCR amplicons (>200bp) spanning the sgRNA target site demonstrated that base editing with optimal reagents virtually eliminates detectable translocations, whereas Cas9 nuclease produces numerous translocations, some at frequencies as high as 1.5%. Notably, larger deletions were also identified at the site of translocation through Sanger sequencing of subcloned junction PCR amplicons (∼500bp) from SpCas9-treated cells (**Supplemental Data 9**). These data suggest the presence of more complex genomic rearrangements similar to those reported previously^9,10^ that are not detected by our current ddPCR assays. Considering the variability in efficiency of nucleic acid delivery between cells by electroporation, it is possible that cells receiving high levels of SpCas9/sgRNA may harbor translocations more frequently.

Although we demonstrate that BE4 substantially reduces DSB induction compared to SpCas9 nuclease and does not produce detectable translocations, the potential remains for undesirable events to occur. For instance, it is possible that the rAPOBEC1 of BE4 could non-specifically edit cytosines in single-stranded DNA during DNA replication^35^. Additionally, the UGIs of BE4 could potentially inhibit uracil DNA glycosylases in a nonspecific fashion, thereby hindering base excision repair of naturally and frequently occurring cytosine deamination in normal mammalian cells^36^. Further studies investigating these potential events should be undertaken prior to clinical translation of base edited cells, though it may be challenging to definitively document such unintended occurrences. Despite these areas of uncertainty, the base editor platform represents a novel approach for highly efficient multiplex engineering of therapeutic primary cells with an improved safety profile compared to current nuclease technologies.

## METHODS

### Cloning & Viral Production

DNA sequences for CD19 chimeric antigen receptor linked by a T2A to RQR8 were synthesized as gBlock Gene Fragments (Integrated DNA Technologies [IDT]). Fragments were Gibson Assembled^37^ into pRRL (www.addgene.org/36247). Gibson reactions were transformed into DH10β E. coli and plated on LB agar with ampicillin. Plasmid DNA was purified from colonies using the GeneJET Plasmid Miniprep Kit (ThermoFisher). Following confirmation by Sanger sequencing, clones were sent to the University of Minnesota Viral Vector & Cloning Core (VVCC) for production and titration of viral particles.

### Guide RNA Design

Guide RNAs (sgRNAs) were designed using the base editing splice-site disruption sgRNA design program SpliceR (https://z.umn.edu/splicer) [Kluesner & Lahr *et al*., *in preparation*]. SpliceR is written in the R statistical programming language (v. 3.4.3). Briefly, SpliceR takes a target Ensembl transcript ID, a base editor PAM variant, and a species as an input. Using the exon and intron sequences from Ensembl, the program extracts the region surrounding every splice site based on a user-specified window. The pattern of N_20_-NGG is then matched to the antisense strand of the extracted sequence. Matched patterns are then scored based on the position of the target motif within the predicted editing window based on previous publications^13^. Subsequently sgRNAs are scored based on their position within the transcript, where sgRNAs earlier in the transcript receive a higher score. pmSTOP inducing gRNAs were designed using the Benchling base editing gRNA design tool (https://benchling.com/pub/liu-base-editor).

### CD3+ T cell Isolation

Peripheral blood mononuclear cells (PBMCs) were isolated from Trima Accel leukoreduction system (LRS) chambers using ammonium chloride-based red blood cell lysis. CD3+ T cells were isolated from the PBMC population by immunomagnetic negative selection using the EasySep Human T cell Isolation Kit (STEMCELL Technologies). T cells were frozen at 10-20×10^6^ cells per 1 mL of Cryostor CS10 (STEMCELL Technologies) and thawed into culture as needed.

### T cell culture

T cells were cultured at 1×10^6^ cells per 1 mL in OpTmizer CTS T cell Expansion SFM containing 2.5% CTS Immune Cell SR (ThermoFisher), L-Glutamine, Penicillin/Streptomycin, N-Acetyl-L-cysteine (10 mM), IL-2 (300 IU), IL-7 (5 ng), and IL-15 (5 ng) at 37°C and 5% CO_2_. T cells were activated with Dynabeads Human T-Activator CD3/CD28 (ThermoFisher) at a 2:1 bead:cell ratio for 48-72 hours prior to electroporation.

### T cell electroporation

After 48 hours, Dynabeads were magnetically removed and cells washed with PBS once prior to resuspension in appropriate electroporation buffer. For singleplex experiments, 3×10^5^ T cells were electroporated with 1 µg of chemically modified sgRNA (Synthego, Menlo Park, CA) and 1.5 µg SpCas9, BE3, or BE4 mRNA (TriLink Biotechnologies) in a 10 µL tip using the Neon Transfection System (ThermoFisher) under the following conditions: 1400 volts, pulse width of 10 milliseconds, 3 pulses. The 4D-Nucleofector (Lonza) and P3 kit was used for multiplex studies with 1×10^6^ T cells per 20 µL cuvette, 1.5-6 µg BE mRNA as indicated, and the Nucleofector program EO-115. RNP were generated by incubation of 10 µg SpCas9 protein (IDT, Coralville Iowa), or 12 µg BE4 protein (Aldevron, Fargo) with 3 µg of chemically modified sgRNA (Synthego) for 15 min at room temperature and electroporated using the Nucleofector program EH-115. T cells were allowed to recover in antibiotic-free medium at 37°C, 5% CO_2_ for 20 minutes following gene transfer, and were then cultured in complete CTS OpTmizer T cell Expansion SFM as described above.

### Lentiviral Transduction

T cells were transduced 24 hours after transfection with pRRL-MND-CAR19-RQR8 lentiviral vector (UMN Viral Vector & Cloning Core) at an MOI of 20 by spinfection on Retronectin (Takara)-coated plates.

### Genomic DNA Analysis

Genomic DNA was isolated from T cells 5 days post-electroporation by spin column-based purification. Base editing efficiency was analyzed on the genomic level by PCR amplification of CRISPR-targeted loci, Sanger sequencing of the PCR amplicons, and subsequent analysis of the Sanger sequencing traces using the web app EditR as previously described (baseeditr.com)^24^. Next generation sequencing (NGS) was also performed on the same PCR amplicons.

### Next Generation Sequencing & Analysis

Primers with Nextera universal primer adaptors (Illumina) were designed to amplify a 375-425 bp site surrounding the region of interest using Primer3Plus. Genomic DNA was PCR-amplified using AccuPrime Taq DNA Polymerase, High Fidelity according to the manufacturer’s protocol (Invitrogen), using the cycle [94°C - 2:00]-30x[94°C - 0:30, 55°C - 0:30, 68°C - 0:30]-[68°C - 5:00]-[4°C - hold]. Amplicons were purified from 1% agarose gel using the QIAquick Gel Extraction Kit (Qiagen). Samples were submitted to the University of Minnesota Genomics Center for subsequent amplification with indexing primers and sequencing on a MiSeq 2×300 bp run (Illumina). A minimum of 1,000 aligned read-pairs were generated per on-target site, and 10,000 read-pairs for off-target sites. Raw fastq files were analyzed against a reference sequence and sgRNA protospacer sequence using the CRISPR/Cas9 editing analysis pipeline CRISPR-DAV as previously described^38^. Output ‘sample_snp.xlsx’ and ‘sample_len.xlsx’ were compiled and analyzed using a custom R markdown script (R v3.4.2).

### Flow Cytometry

Prior to flow cytometry, singleplex PDCD1 disrupted T cells were re-stimulated using CD3/CD28 Dynabeads for 48 hours as described above. In multiplex experiments with TRAC knockout, T cells were activated with Phorbol 12-myristate 13-acetate (PMA; 100 ng/mL; Sigma-Aldrich) and ionomycin (250 ng/mL; MilliporeSigma) for 24 hours. T cells treated with PMA/ionomycin were washed with PBS, resuspended in culture medium, and incubated for an additional 24 hours prior to flow cytometry. T cells were stained with fluorophore-conjugated anti-human CD3 (BD Biosciences), B2M (BioLegend), and CD279 (PD-1) (BioLegend) antibodies. Anti-human CD34 monoclonal antibody (QBEnd10) (ThermoFisher) was used to detect CD19-T2A-RQR8 CAR expression. Fixable Viability Dye eFluor 780 or LIVE/DEAD Fixable Aqua Dead Cell Stain (ThermoFisher) were used to assess cell viability. T cells were acquired on LSR II or LSRFortessa flow cytometers using FACSDiva software, and data were analyzed using FlowJo v10 software. As stimulation does not uniformly upregulate PD-1 expression in all control T cells, PD-1^+^ cell frequencies were normalized. The ratio (*r*_PD1_) of PD-1^+^ cells to PD-1^-^ subpopulations in control samples was used to calculate the normalized values (*F’*_pos_’ and *F’*_neg_’) of PD-1^+^ and PD-1^-^ subpopulations from the non-normalized values (F°_pos_ and F°_pos_) for all samples as follows:

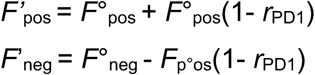

### Translocation assay

Translocation PCR assays were designed using PrimerQuest software (Integrated DNA Technologies, Coralville IA) using settings for 2 primers+probe qPCR. Each sample was run as a duplexed assay consisting of an internal reference primer+probe set (HEX) and an experimental primer+probe set (FAM). Primers and probes were ordered from IDT. Reactions were set up using the ddPCR Supermix for Probes (no dUTP) (Biorad, Hercules, CA) with 200ng of genomic DNA per assay according to manufacturer instructions. Droplets were generated and analyzed using QX200 Droplet-digital PCR system (Bio-Rad).

### Cytotoxicity assay

Luciferase-expressing K562, Raji, or Raji-PDL1 cells were seeded into a 96-well round-bottom plate (3×10^4^ cells/well). T cells were counted and added to the wells in triplicate at the indicated E:T ratios. Target cells without effectors served as a negative control (spontaneous cell death) and target cells incubated with 1% NP-40 served as positive control (maximum killing). Co-cultures were incubated at 37°C for 48 hours. After incubation, D-luciferin (potassium salt; Gold Biotechnology) was added to each well at a final concentration of 25 µg/mL and incubated 10 minutes before imaging. Luminescence was read in endpoint mode using BioTek Synergy microplate reader. Target cells with no effectors were set as 100% survival and killing in experimental samples was measured against this baseline.

### Immunoblotting assay

Proteins were isolated from 1×10^6^ cells in complete RIPA buffer with protease and phosphatase inhibitors (Sigma-Aldrich, COEDTAF-RO, P5726, and P0044). Total protein was quantified using the Pierce BCA Protein Assay Kit (Thermo Fisher Scientific Inc., 23225) according to the manufacturer’s protocol. Protein samples were run and analyzed on the Wes platform after being denatured at 95°C for 5 minutes according to the manufacturer’s protocol (ProteinSimple). Primary antibodies against SpCas9 (Cell Signaling, #14697) and actin (Cell Signaling, #8457) were used at 1:100 and 1:50 dilutions, respectively, in kit-supplied buffer and platform-optimized secondary antibodies were purchased from Protein Simple.

### Data analysis and visualization

All statistical analyses were performed in R studio. The level of significance was set at *α* = 0.05. Data were subjected to analyses for the assumptions of normality and homeodascity prior to statistical testing. Student’s pairwise one-tailed or two-tailed t-tests were used as indicated in the text. Data were visualized using either Prism 8 (Graphpad), or R studio employing various tidyverse (www.tidyverse.org/) and Bioconductor (www.bioconductor.org/) packages.

### Data availability

Next-generation sequencing reads will be deposited in the NCBI Sequence Read Archive database prior to publication.

## Supporting information

## ACKNOWLEDGMENTS

This works was supported by grants from the Children’s Cancer Research fund, the Sobiech Osteosarcoma Fund Award, and the University of Minnesota Department of Pediatrics. B.R.W was supported by the Emerging Scientist award from the Children’s Cancer Research Fund. We thank Kenny Beckman and John Garbe from the University of Minnesota Genomics Center for advice on performing NGS; Madison Vignes and Lindsey Sumstad from the University of Minnesota for assistance in preparing NGS samples; Jason Gehrke and Darrell Johnson for advice on ddPCR assays; and Ezequiel Marron at the University of Minnesota Viral Vector & Cloning Core for producing lentivirus.

## AUTHOR CONTRIBUTIONS

B.R.W conceived of the project, designed experiments, and performed data analysis with assistance from C.L. C.L. performed all experiments with assistance from B.R.W. and M.D.D. Guide RNA design was carried out by M.G.K and W.S.L with assistance from B.R.W. Preparation of samples for NGS was carried out by M.G.K, W.S.L., and G.D. M.G.K analyzed NGS data. Western blot analysis was performed by N.J.S. using recombinant BE3 protein produced by K.N.L. and W.R.G. Translocation PCR and sequencing was carried out by A.M. with assistance from M.J.O. Multiplex flow cytometry analysis and cytokine functionality assays were carried out by M.J.J. with assistance from C.L. Cytotoxicity assays were carried out by E.J.P. B.S.M. conceived of the project, designed experiments, and directed the research with assistance from B.R.W. B.S.M. and B.R.W wrote the manuscript with input from all the authors.

## COMPETEING INTERESTS

B.R.W. and B.S.M. are consultants for Beam Therapeutics. B.R.W and B.S.M. have financial interests in Beam Therapeutics. Both B.R.W. and B.S.M.’s interests were reviewed and are managed by the University of Minnesota in accordance with their conflict of interest policies.

