## Supplementary material for "Highly efficient multiplex human T cell engineering without double-strand breaks using Cas9 base editors"

**Supplemental Table 1. Single-guide RNA (sgRNA) information.**

| Gene | gRNA Name | 5' - gRNA Sequence - 3' | Orientation | Target base(s) | Predicted Outcome |
|---|---|---|---|---|---|
| *PDCD1* | Ex. 1 SD | CACCTACCTAAGAACCATCC | Antisense | C7 | Splice donor disruption: GT → AT |
| *PDCD1* | Ex. 2 SA | GGAGTCTGAGAGATGGAGAG | Antisense | C6 | Splice acceptor disruption: AG → AA |
| *PDCD1* | Ex. 3 SA | TTCTCTCTGGAAGGGCACAA | Antisense | C7 | Splice acceptor disruption: AG → AA |
| *PDCD1* | Ex. 3 SD | GACGTTACCTCGTGCGGCCC | Antisense | C8 | Splice donor disruption: GT → AT |
| *PDCD1* | Ex. 4 SA | CCTGCAGAGAAACACACTTG | Antisense | C2 | Splice acceptor disruption: AG → AA |
| *PDCD1* | Ex. 2 pmSTOP | GGGGTTCCAGGGCCTGTCTG | Antisense | C7, C8 | pmSTOP induction: TGG (Trp) → TAG, TGA, TAA |
| *PDCD1* | Ex. 3 pmSTOP_1 | CAGTTCCAAACCCTGGTGGT | Sense | C7 | pmSTOP induction: CAA (Gln) → TAA |
| *PDCD1* | Ex. 3 pmSTOP_2 | GGACCCAGACTAGCAGCACC | Antisense | C5, C6 | pmSTOP induction: TGG (Trp) → TAG, TGA, TAA |
| *TRAC* | Ex. 1 SD | CTTACCTGGGCTGGGGAAGA | Antisense | C5 | Splice donor disruption: GT → AT |
| *TRAC* | Ex. 3 SA | TTCGTATCTGTAAAACCAAG | Antisense | C8 | Splice acceptor disruption: AG → AA |
| *TRAC* | Ex. 3 pmSTOP_1 | TTTCAAAACCTGTCAGTGAT | Sense | C4 | pmSTOP induction: CAA (Gln) → TAA |
| *TRAC* | Ex. 3 pmSTOP_2 | TTCAAAACCTGTCAGTGATT | Sense | C3 | pmSTOP induction: CAA (Gln) → TAA |
| *B2M* | Ex. 1 SD | ACTCACGCTGGATAGCCTCC | Antisense | C6 | Splice donor disruption: GT → AT |
| *B2M* | Ex. 3 SA | TCGATCTATGAAAAAGACAG | Antisense | C6 | Splice acceptor disruption: AG → AA |
| *B2M* | Ex. 2 pmSTOP | CTTACCCCACTTAACTATCT | Antisense | C7, C8 | pmSTOP induction: TGG (Trp) → TAG, TGA, TAA |

### Supplemental Table 2. Computationally predicted candidate off-target sites.

| Site Name | Primer Name | Primer Sequence | Off-Target Sequence | Alignment | Gene | Coordinates |
|---|---|---|---|---|---|---|
| B2M_Ex1_SD_OnT | B2M_Ex1_SD_OnT Fwd 1 | TCGTCGGCAGCGTCAGATGTGTATAAGAGACAGATCCAGCCCTGGACTAGC | ACTCAcGCTGGATAGCCTCC | .................... | B2M | chr15:45003795-45003817 |
| B2M_Ex1_SD_OnT | B2M_Ex1_SD_OnT Rev 1 | GTCTCGTGGGCTCGGAGATGTGTATAAGAGACAGCTCTCTCTAACCTGGCACTG | ACTCAcGCTGGATAGCCTCC | .................... | B2M | chr15:45003795-45003817 |
| B2M_Ex1_SD_OnT | B2M_Ex1_SD_OnT Fwd 2 | TCGTCGGCAGCGTCAGATGTGTATAAGAGACAGATCCAGCCCTGGACTAGC | ACTCAcGCTGGATAGCCTCC | .................... | B2M | chr15:45003795-45003817 |
| B2M_Ex1_SD_OnT | B2M_Ex1_SD_OnT Rev 2 | GTCTCGTGGGCTCGGAGATGTGTATAAGAGACAGCCTCTCTCTAACCTGGCACT | ACTCAcGCTGGATAGCCTCC | .................... | B2M | chr15:45003795-45003817 |
| B2M_Ex1_SD_OT1 | B2M_Ex1_SD_OT1 Fwd 1 | TCGTCGGCAGCGTCAGATGTGTATAAGAGACAGCCCAGACATGAGAAGGTTAT | TCTGCCCCTGGATAGCCTCC | T..GC.C............. | PDE11A | chr2:178777091-178777113 |
| B2M_Ex1_SD_OT1 | B2M_Ex1_SD_OT1 Rev 1 | GTCTCGTGGGCTCGGAGATGTGTATAAGAGACAGCTTTACAGGCTCCCCTTC | TCTGCCCCTGGATAGCCTCC | T..GC.C............. | PDE11A | chr2:178777091-178777113 |
| B2M_Ex1_SD_OT1 | B2M_Ex1_SD_OT1 Fwd 2 | TCGTCGGCAGCGTCAGATGTGTATAAGAGACAGCCCAGACATGAGAAGGTTAT | TCTGCCCCTGGATAGCCTCC | T..GC.C............. | PDE11A | chr2:178777091-178777113 |
| B2M_Ex1_SD_OT1 | B2M_Ex1_SD_OT1 Rev 2 | GTCTCGTGGGCTCGGAGATGTGTATAAGAGACAGGTCTAGGCGTCCTCATCAC | TCTGCCCCTGGATAGCCTCC | T..GC.C............. | PDE11A | chr2:178777091-178777113 |
| B2M_Ex1_SD_OT2 | B2M_Ex1_SD_OT2 Fwd 1 | TCGTCGGCAGCGTCAGATGTGTATAAGAGACAGGGATGTTCTTTGGTGTTTGT | ACTCACCTTCCATAGCCTCC | ......CT.CC......... | ZNF519 | chr18:14090054-14090076 |
| B2M_Ex1_SD_OT2 | B2M_Ex1_SD_OT2 Rev 1 | GTCTCGTGGGCTCGGAGATGTGTATAAGAGACAGGACTCCGTCTCTGAACACTC | ACTCACCTTCCATAGCCTCC | ......CT.CC......... | ZNF519 | chr18:14090054-14090076 |
| B2M_Ex1_SD_OT2 | B2M_Ex1_SD_OT2 Fwd 2 | TCGTCGGCAGCGTCAGATGTGTATAAGAGACAGGAATGTTGGGATGTTCTTTG | ACTCACCTTCCATAGCCTCC | ......CT.CC......... | ZNF519 | chr18:14090054-14090076 |
| B2M_Ex1_SD_OT2 | B2M_Ex1_SD_OT2 Rev 2 | GTCTCGTGGGCTCGGAGATGTGTATAAGAGACAGGACTCCGTCTCTGAACACTC | ACTCACCTTCCATAGCCTCC | ......CT.CC......... | ZNF519 | chr18:14090054-14090076 |
| B2M_Ex1_SD_OT3 | B2M_Ex1_SD_OT3 Fwd 1 | TCGTCGGCAGCGTCAGATGTGTATAAGAGACAGATAGTTGCCATTTCTGCTTG | GCTCCTGCTGCATAGCCTCC | G...CT....C......... | KLF13 | chr15:31648182-31648204 |
| B2M_Ex1_SD_OT3 | B2M_Ex1_SD_OT3 Rev 1 | GTCTCGTGGGCTCGGAGATGTGTATAAGAGACAGGAGGGTGAAAGACTGAAAAA | GCTCCTGCTGCATAGCCTCC | G...CT....C......... | KLF13 | chr15:31648182-31648204 |
| B2M_Ex1_SD_OT3 | B2M_Ex1_SD_OT3 Fwd 2 | TCGTCGGCAGCGTCAGATGTGTATAAGAGACAGAATAGTTGCCATTTCTGCTT | GCTCCTGCTGCATAGCCTCC | G...CT....C......... | KLF13 | chr15:31648182-31648204 |
| B2M_Ex1_SD_OT3 | B2M_Ex1_SD_OT3 Rev 2 | GTCTCGTGGGCTCGGAGATGTGTATAAGAGACAGGAGGGTGAAAGACTGAAAAA | GCTCCTGCTGCATAGCCTCC | G...CT....C......... | KLF13 | chr15:31648182-31648204 |
| B2M_Ex1_SD_OT4 | B2M_Ex1_SD_OT4 Fwd 1 | TCGTCGGCAGCGTCAGATGTGTATAAGAGACAGTCTTTTTGTGAAGGCTTTTC | TCTCACTGTGGTTAGCCTCC | T.....TG...T........ | NA | chr11:123367986-123368008 |
| B2M_Ex1_SD_OT4 | B2M_Ex1_SD_OT4 Rev 1 | GTCTCGTGGGCTCGGAGATGTGTATAAGAGACAGTTTGTCACCAGCTGATTGTA | TCTCACTGTGGTTAGCCTCC | T.....TG...T........ | NA | chr11:123367986-123368008 |
| B2M_Ex1_SD_OT4 | B2M_Ex1_SD_OT4 Fwd 2 | TCGTCGGCAGCGTCAGATGTGTATAAGAGACAGTCTTTTTGTGAAGGCTTTTC | TCTCACTGTGGTTAGCCTCC | T.....TG...T........ | NA | chr11:123367986-123368008 |
| B2M_Ex1_SD_OT4 | B2M_Ex1_SD_OT4 Rev 2 | GTCTCGTGGGCTCGGAGATGTGTATAAGAGACAGTGTCACCAGCTGATTGTAGA | TCTCACTGTGGTTAGCCTCC | T.....TG...T........ | NA | chr11:123367986-123368008 |
| B2M_Ex1_SD_OT5 | B2M_Ex1_SD_OT5 Fwd 1 | TCGTCGGCAGCGTCAGATGTGTATAAGAGACAGCTCAAGAGCAAGAAACCAGT | TCTCTCACTGGATAGCCTAC | T...T.A...........A. | NA | chr1:144373808-144373830 |
| B2M_Ex1_SD_OT5 | B2M_Ex1_SD_OT5 Rev 1 | GTCTCGTGGGCTCGGAGATGTGTATAAGAGACAGAGGACTTTGAAATACCAGCA | TCTCTCACTGGATAGCCTAC | T...T.A...........A. | NA | chr1:144373808-144373830 |
| B2M_Ex1_SD_OT5 | B2M_Ex1_SD_OT5 Fwd 2 | TCGTCGGCAGCGTCAGATGTGTATAAGAGACAGGTGAGTCAGCAGCTCAAGA | TCTCTCACTGGATAGCCTAC | T...T.A...........A. | NA | chr1:144373808-144373830 |
| B2M_Ex1_SD_OT5 | B2M_Ex1_SD_OT5 Rev 2 | GTCTCGTGGGCTCGGAGATGTGTATAAGAGACAGAGGACTTTGAAATACCAGCA | TCTCTCACTGGATAGCCTAC | T...T.A...........A. | NA | chr1:144373808-144373830 |
| B2M_Ex1_SD_OT6 | B2M_Ex1_SD_OT6 Fwd 1 | TCGTCGGCAGCGTCAGATGTGTATAAGAGACAGCTCAAGAGCAAGAAACCAGT | TCTCTCACTGGATAGCCTAC | T...T.A...........A. | LINC01138 | chr1:147964949-147964971 |
| B2M_Ex1_SD_OT6 | B2M_Ex1_SD_OT6 Rev 1 | GTCTCGTGGGCTCGGAGATGTGTATAAGAGACAGAGGACTTTGAAATACCAGCA | TCTCTCACTGGATAGCCTAC | T...T.A...........A. | LINC01138 | chr1:147964949-147964971 |
| B2M_Ex1_SD_OT6 | B2M_Ex1_SD_OT6 Fwd 2 | TCGTCGGCAGCGTCAGATGTGTATAAGAGACAGGTGAGTCAGCAGCTCAAGA | TCTCTCACTGGATAGCCTAC | T...T.A...........A. | LINC01138 | chr1:147964949-147964971 |
| B2M_Ex1_SD_OT6 | B2M_Ex1_SD_OT6 Rev 2 | GTCTCGTGGGCTCGGAGATGTGTATAAGAGACAGAGGACTTTGAAATACCAGCA | TCTCTCACTGGATAGCCTAC | T...T.A...........A. | LINC01138 | chr1:147964949-147964971 |
| B2M_Ex1_SD_OT7 | B2M_Ex1_SD_OT7 Fwd 1 | TCGTCGGCAGCGTCAGATGTGTATAAGAGACAGCTCAAGAGCAAGAAACCAGT | TCTCTCACTGGATAGCCTAC | T...T.A...........A. | NA | chr1:149543388-149543410 |
| B2M_Ex1_SD_OT7 | B2M_Ex1_SD_OT7 Rev 1 | GTCTCGTGGGCTCGGAGATGTGTATAAGAGACAGAGGACTTTGAAATACCAGCA | TCTCTCACTGGATAGCCTAC | T...T.A...........A. | NA | chr1:149543388-149543410 |
| B2M_Ex1_SD_OT7 | B2M_Ex1_SD_OT7 Fwd 2 | TCGTCGGCAGCGTCAGATGTGTATAAGAGACAGGTGAGTCAGCAGCTCAAGA | TCTCTCACTGGATAGCCTAC | T...T.A...........A. | NA | chr1:149543388-149543410 |
| B2M_Ex1_SD_OT7 | B2M_Ex1_SD_OT7 Rev 2 | GTCTCGTGGGCTCGGAGATGTGTATAAGAGACAGAGGACTTTGAAATACCAGCA | TCTCTCACTGGATAGCCTAC | T...T.A...........A. | NA | chr1:149543388-149543410 |
| B2M_Ex1_SD_OT8 | B2M_Ex1_SD_OT8 Fwd 1 | TCGTCGGCAGCGTCAGATGTGTATAAGAGACAGAGCTCAGCTTGCTCCACT | AGCCACCCTGGAGAGCCTCC | .GC...C....G........ | LRRC8E | chr19:7964982-7965004 |
| B2M_Ex1_SD_OT8 | B2M_Ex1_SD_OT8 Rev 1 | GTCTCGTGGGCTCGGAGATGTGTATAAGAGACAGGCTTCTTGAGGCTGTTCA | AGCCACCCTGGAGAGCCTCC | .GC...C....G........ | LRRC8E | chr19:7964982-7965004 |
| B2M_Ex1_SD_OT8 | B2M_Ex1_SD_OT8 Fwd 2 | TCGTCGGCAGCGTCAGATGTGTATAAGAGACAGAGCTCAGCTTGCTCCACT | AGCCACCCTGGAGAGCCTCC | .GC...C....G........ | LRRC8E | chr19:7964982-7965004 |
| B2M_Ex1_SD_OT8 | B2M_Ex1_SD_OT8 Rev 2 | GTCTCGTGGGCTCGGAGATGTGTATAAGAGACAGCTTCTTGAGGCTGTTCAGG | AGCCACCCTGGAGAGCCTCC | .GC...C....G........ | LRRC8E | chr19:7964982-7965004 |
| B2M_Ex1_SD_OT9 | B2M_Ex1_SD_OT9 Fwd 1 | TCGTCGGCAGCGTCAGATGTGTATAAGAGACAGCTTTCGCCAGCTCAAAAA | ACTCCCGCTGGAAAGCCTGC | ....C.......A.....G. | NA | chr17:20755870-20755892 |
| B2M_Ex1_SD_OT9 | B2M_Ex1_SD_OT9 Rev 1 | GTCTCGTGGGCTCGGAGATGTGTATAAGAGACAGCAGCGAGCTCTACTGGTG | ACTCCCGCTGGAAAGCCTGC | ....C.......A.....G. | NA | chr17:20755870-20755892 |
| B2M_Ex1_SD_OT9 | B2M_Ex1_SD_OT9 Fwd 2 | TCGTCGGCAGCGTCAGATGTGTATAAGAGACAGCTTTCGCCAGCTCAAAAA | ACTCCCGCTGGAAAGCCTGC | ....C.......A.....G. | NA | chr17:20755870-20755892 |
| B2M_Ex1_SD_OT9 | B2M_Ex1_SD_OT9 Rev 2 | GTCTCGTGGGCTCGGAGATGTGTATAAGAGACAGCGAGCTCTACTGGTGCTG | ACTCCCGCTGGAAAGCCTGC | ....C.......A.....G. | NA | chr17:20755870-20755892 |
| B2M_Ex1_SD_OT10 | B2M_Ex1_SD_OT10 Fwd 1 | TCGTCGGCAGCGTCAGATGTGTATAAGAGACAGGGTGGATTACAATGGAAACA | CCTCCCGCTGTGTAGCCTCC | C...C.....TG........ | COL13A1 | chr10:71685789-71685811 |
| B2M_Ex1_SD_OT10 | B2M_Ex1_SD_OT10 Rev 1 | GTCTCGTGGGCTCGGAGATGTGTATAAGAGACAGCATGAAAGGGGGTTATACAT | CCTCCCGCTGTGTAGCCTCC | C...C.....TG........ | COL13A1 | chr10:71685789-71685811 |
| B2M_Ex1_SD_OT10 | B2M_Ex1_SD_OT10 Fwd 2 | TCGTCGGCAGCGTCAGATGTGTATAAGAGACAGGGATTACAATGGAAACATCAA | CCTCCCGCTGTGTAGCCTCC | C...C.....TG........ | COL13A1 | chr10:71685789-71685811 |
| B2M_Ex1_SD_OT10 | B2M_Ex1_SD_OT10 Rev 2 | GTCTCGTGGGCTCGGAGATGTGTATAAGAGACAGCATGAAAGGGGGTTATACAT | CCTCCCGCTGTGTAGCCTCC | C...C.....TG........ | COL13A1 | chr10:71685789-71685811 |
| PD-1_Ex_SD_OnT | PD-1_Ex.1_SD_OnT Fwd 1 | TCGTCGGCAGCGTCAGATGTGTATAAGAGACAGCTGCCAGGGACTGAGAGT | CACCTAcCTAAGAACCATCC | .................... | PDCD1 | chr2:242800908-242800930 |
| PD-1_Ex_SD_OnT | PD-1_Ex.1_SD_OnT Rev 1 | GTCTCGTGGGCTCGGAGATGTGTATAAGAGACAGGTGGATGTGGAGGAAGAG | CACCTAcCTAAGAACCATCC | .................... | PDCD1 | chr2:242800908-242800930 |
| PD-1_Ex_SD_OnT | PD-1_Ex.1_SD_OnT Fwd 2 | TCGTCGGCAGCGTCAGATGTGTATAAGAGACAGCTGCCAGGGACTGAGAGT | CACCTAcCTAAGAACCATCC | .................... | PDCD1 | chr2:242800908-242800930 |
| PD-1_Ex_SD_OnT | PD-1_Ex.1_SD_OnT Rev 2 | GTCTCGTGGGCTCGGAGATGTGTATAAGAGACAGACGTGGATGTGGAGGAAG | CACCTAcCTAAGAACCATCC | .................... | PDCD1 | chr2:242800908-242800930 |
| PD-1_Ex_SD_OT1 | PD-1_Ex.1_SD_OT1 Fwd 1 | TCGTCGGCAGCGTCAGATGTGTATAAGAGACAGTTTCACTTCTATCCCACACC | CGGCCACCTGAGAACCATCC | .GG.C....G.......... | CDKL5 | chrX:18663622-18663644 |
| PD-1_Ex_SD_OT1 | PD-1_Ex.1_SD_OT1 Rev 1 | GTCTCGTGGGCTCGGAGATGTGTATAAGAGACAGAAAGTTCTCTGGTTCCTGTG | CGGCCACCTGAGAACCATCC | .GG.C....G.......... | CDKL5 | chrX:18663622-18663644 |
| PD-1_Ex_SD_OT1 | PD-1_Ex.1_SD_OT1 Fwd 2 | TCGTCGGCAGCGTCAGATGTGTATAAGAGACAGTTTCACTTCTATCCCACACC | CGGCCACCTGAGAACCATCC | .GG.C....G.......... | CDKL5 | chrX:18663622-18663644 |
| PD-1_Ex_SD_OT1 | PD-1_Ex.1_SD_OT1 Rev 2 | GTCTCGTGGGCTCGGAGATGTGTATAAGAGACAGGAAAAGTTCTCTGGTTCCTG | CGGCCACCTGAGAACCATCC | .GG.C....G.......... | CDKL5 | chrX:18663622-18663644 |
| PD-1_Ex_SD_OT2 | PD-1_Ex.1_SD_OT2 Fwd 1 | TCGTCGGCAGCGTCAGATGTGTATAAGAGACAGGGGCCACTTGTGCTAGAG | ACCTGACCTAAGAACCATCC | AC.TG............... | ST8SIA2 | chr15:92989355-92989377 |
| PD-1_Ex_SD_OT2 | PD-1_Ex.1_SD_OT2 Rev 1 | GTCTCGTGGGCTCGGAGATGTGTATAAGAGACAGACTAGTGCCCATGATAGCAG | ACCTGACCTAAGAACCATCC | AC.TG............... | ST8SIA2 | chr15:92989355-92989377 |
| PD-1_Ex_SD_OT2 | PD-1_Ex.1_SD_OT2 Fwd 2 | TCGTCGGCAGCGTCAGATGTGTATAAGAGACAGCATGTACACGTCTGACCACT | ACCTGACCTAAGAACCATCC | AC.TG............... | ST8SIA2 | chr15:92989355-92989377 |
| PD-1_Ex_SD_OT2 | PD-1_Ex.1_SD_OT2 Rev 2 | GTCTCGTGGGCTCGGAGATGTGTATAAGAGACAGGAATTTACTAGTGCCCATGAT | ACCTGACCTAAGAACCATCC | AC.TG............... | ST8SIA2 | chr15:92989355-92989377 |
| PD-1_Ex_SD_OT3 | PD-1_Ex.1_SD_OT3 Fwd 1 | TCGTCGGCAGCGTCAGATGTGTATAAGAGACAGACACTAACGATTGCTGATGA | CTTCTATCTCAGAACCATCC | .TT...T..C.......... | QRFPR | chr4:122259478-122259500 |
| PD-1_Ex_SD_OT3 | PD-1_Ex.1_SD_OT3 Rev 1 | GTCTCGTGGGCTCGGAGATGTGTATAAGAGACAGTTTCTCACTCGCTCTTTCTC | CTTCTATCTCAGAACCATCC | .TT...T..C.......... | QRFPR | chr4:122259478-122259500 |
| PD-1_Ex_SD_OT3 | PD-1_Ex.1_SD_OT3 Fwd 2 | TCGTCGGCAGCGTCAGATGTGTATAAGAGACAGCACTAACGATTGCTGATGAC | CTTCTATCTCAGAACCATCC | .TT...T..C.......... | QRFPR | chr4:122259478-122259500 |
| PD-1_Ex_SD_OT3 | PD-1_Ex.1_SD_OT3 Rev 2 | GTCTCGTGGGCTCGGAGATGTGTATAAGAGACAGTTTCTCACTCGCTCTTTCTC | CTTCTATCTCAGAACCATCC | .TT...T..C.......... | QRFPR | chr4:122259478-122259500 |
| PD-1_Ex_SD_OT4 | PD-1_Ex.1_SD_OT4 Fwd 1 | TCGTCGGCAGCGTCAGATGTGTATAAGAGACAGTTTCTAGCTTCTGCCTTCTC | TACCCAGCTCAGAACCATCC | T...C.G..C.......... | NA | chr16:8315346-8315368 |
| PD-1_Ex_SD_OT4 | PD-1_Ex.1_SD_OT4 Rev 1 | GTCTCGTGGGCTCGGAGATGTGTATAAGAGACAGTGCTTTTCAGAATTGATGTG | TACCCAGCTCAGAACCATCC | T...C.G..C.......... | NA | chr16:8315346-8315368 |
| PD-1_Ex_SD_OT4 | PD-1_Ex.1_SD_OT4 Fwd 2 | TCGTCGGCAGCGTCAGATGTGTATAAGAGACAGATTTCTAGCTTCTGCCTTCTC | TACCCAGCTCAGAACCATCC | T...C.G..C.......... | NA | chr16:8315346-8315368 |
| PD-1_Ex_SD_OT4 | PD-1_Ex.1_SD_OT4 Rev 2 | GTCTCGTGGGCTCGGAGATGTGTATAAGAGACAGTGCTTTTCAGAATTGATGTG | TACCCAGCTCAGAACCATCC | T...C.G..C.......... | NA | chr16:8315346-8315368 |
| PD-1_Ex_SD_OT5 | PD-1_Ex.1_SD_OT5 Fwd 1 | TCGTCGGCAGCGTCAGATGTGTATAAGAGACAGTGTAGTTCAGGGCTGTTAGG | CACTCACTTAAGTACCATCC | ...TC..T....T....... | NA | chr3:148663173-148663195 |
| PD-1_Ex_SD_OT5 | PD-1_Ex.1_SD_OT5 Rev 1 | GTCTCGTGGGCTCGGAGATGTGTATAAGAGACAGTTTCAAAACTAACCAATCTGC | CACTCACTTAAGTACCATCC | ...TC..T....T....... | NA | chr3:148663173-148663195 |
| PD-1_Ex_SD_OT5 | PD-1_Ex.1_SD_OT5 Fwd 2 | TCGTCGGCAGCGTCAGATGTGTATAAGAGACAGGCTGTTAGGGAGACTGAGAA | CACTCACTTAAGTACCATCC | ...TC..T....T....... | NA | chr3:148663173-148663195 |
| PD-1_Ex_SD_OT5 | PD-1_Ex.1_SD_OT5 Rev 2 | GTCTCGTGGGCTCGGAGATGTGTATAAGAGACAGTTTCAAAACTAACCAATCTGC | CACTCACTTAAGTACCATCC | ...TC..T....T....... | NA | chr3:148663173-148663195 |
| PD-1_Ex_SD_OT6 | PD-1_Ex.1_SD_OT6 Fwd 1 | TCGTCGGCAGCGTCAGATGTGTATAAGAGACAGGTGAGGAGGCATCCGAGT | CACAAACCTGAGAACCATCG | ...AA....G.........G | ATP11A | chr13:113377068-113377090 |
| PD-1_Ex_SD_OT6 | PD-1_Ex.1_SD_OT6 Rev 1 | GTCTCGTGGGCTCGGAGATGTGTATAAGAGACAGGCAAATTAAAGAATCTCTGAAAA | CACAAACCTGAGAACCATCG | ...AA....G.........G | ATP11A | chr13:113377068-113377090 |
| PD-1_Ex_SD_OT6 | PD-1_Ex.1_SD_OT6 Fwd 2 | TCGTCGGCAGCGTCAGATGTGTATAAGAGACAGGAAGGAGGGGGTGAGGAG | CACAAACCTGAGAACCATCG | ...AA....G.........G | ATP11A | chr13:113377068-113377090 |
| PD-1_Ex_SD_OT6 | PD-1_Ex.1_SD_OT6 Rev 2 | GTCTCGTGGGCTCGGAGATGTGTATAAGAGACAGAAAGTAAAGCATTTCTGAATCC | CACAAACCTGAGAACCATCG | ...AA....G.........G | ATP11A | chr13:113377068-113377090 |
| PD-1_Ex_SD_OT7 | PD-1_Ex.1_SD_OT7 Fwd 1 | TCGTCGGCAGCGTCAGATGTGTATAAGAGACAGACCCTGCACAGAACCTATAA | CACCTCCATTTGAACCATCC | .....C.A.TT......... | GRID1 | chr10:87654974-87654996 |

### Supplemental Table 2 continued. Computationally predicted candidate off-target sites.

| | | | | | | |
|---|---|---|---|---|---|---|
| PD-1_Ex_SD_OT7 | PD-1_Ex.1_SD_OT7 Rev 1 | GTCTCGTGGGCTCGGAGATGTGTATAAGAGACAGAAGGACTTGGCTTGTCTTCT | CACCTCCATTTGAACCATCC | .....C.A.TT......... | GRID1 | chr10:87654974-87654996 |
| PD-1_Ex_SD_OT7 | PD-1_Ex.1_SD_OT7 Fwd 2 | TCGTCGGCAGCGTCAGATGTGTATAAGAGACAGACCCTGCACAGAACCTATAA | CACCTCCATTTGAACCATCC | .....C.A.TT......... | GRID1 | chr10:87654974-87654996 |
| PD-1_Ex_SD_OT7 | PD-1_Ex.1_SD_OT7 Rev 2 | GTCTCGTGGGCTCGGAGATGTGTATAAGAGACAGGACTTGGCTTGTCTTCTGAT | CACCTCCATTTGAACCATCC | .....C.A.TT......... | GRID1 | chr10:87654974-87654996 |
| PD-1_Ex_SD_OT8 | PD-1_Ex.1_SD_OT8 Fwd 1 | TCGTCGGCAGCGTCAGATGTGTATAAGAGACAGCCTATTTCATATTGGGTGGA | CACCCACCTAAGCACCATCT | ....C.......C......T | NA | chr6:11660928-11660950 |
| PD-1_Ex_SD_OT8 | PD-1_Ex.1_SD_OT8 Rev 1 | GTCTCGTGGGCTCGGAGATGTGTATAAGAGACAGTTCAAAACACAGGGAAAACT | CACCCACCTAAGCACCATCT | ....C.......C......T | NA | chr6:11660928-11660950 |
| PD-1_Ex_SD_OT8 | PD-1_Ex.1_SD_OT8 Fwd 2 | TCGTCGGCAGCGTCAGATGTGTATAAGAGACAGTTTCATATTGGGTGGATTGT | CACCCACCTAAGCACCATCT | ....C.......C......T | NA | chr6:11660928-11660950 |
| PD-1_Ex_SD_OT8 | PD-1_Ex.1_SD_OT8 Rev 2 | GTCTCGTGGGCTCGGAGATGTGTATAAGAGACAGTTCAAAACACAGGGAAAACT | CACCCACCTAAGCACCATCT | ....C.......C......T | NA | chr6:11660928-11660950 |
| PD-1_Ex_SD_OT9 | PD-1_Ex.1_SD_OT9 Fwd 1 | TCGTCGGCAGCGTCAGATGTGTATAAGAGACAGGCAGCATCACCTGTGTAAC | CACCTTCATCAGAACCATCT | .....T.A.C.........T | NA | chr1:154444930-154444952 |
| PD-1_Ex_SD_OT9 | PD-1_Ex.1_SD_OT9 Rev 1 | GTCTCGTGGGCTCGGAGATGTGTATAAGAGACAGGAAGTAAAAAGCAGAGGAAGC | CACCTTCATCAGAACCATCT | .....T.A.C.........T | NA | chr1:154444930-154444952 |
| PD-1_Ex_SD_OT9 | PD-1_Ex.1_SD_OT9 Fwd 2 | TCGTCGGCAGCGTCAGATGTGTATAAGAGACAGAGCAGCATCACCTGTGTAA | CACCTTCATCAGAACCATCT | .....T.A.C.........T | NA | chr1:154444930-154444952 |
| PD-1_Ex_SD_OT9 | PD-1_Ex.1_SD_OT9 Rev 2 | GTCTCGTGGGCTCGGAGATGTGTATAAGAGACAGGAAGTAAAAAGCAGAGGAAGC | CACCTTCATCAGAACCATCT | .....T.A.C.........T | NA | chr1:154444930-154444952 |
| PD-1_Ex_SD_OT10 | PD-1_Ex.1_SD_OT10 Fwd 1 | TCGTCGGCAGCGTCAGATGTGTATAAGAGACAGTACATGTTTTCTATATGAGGCATT | CAGCTATCTCAGAACCTTCC | ..G...T..C......T... | NA | chr5:66830623-66830645 |
| PD-1_Ex_SD_OT10 | PD-1_Ex.1_SD_OT10 Rev 1 | GTCTCGTGGGCTCGGAGATGTGTATAAGAGACAGCTTCCTCTAAGTCTCAGCTCAT | CAGCTATCTCAGAACCTTCC | ..G...T..C......T... | NA | chr5:66830623-66830645 |
| PD-1_Ex_SD_OT10 | PD-1_Ex.1_SD_OT10 Fwd 2 | TCGTCGGCAGCGTCAGATGTGTATAAGAGACAGTTTTCTATTTTTCCATACTTTTTAG | CAGCTATCTCAGAACCTTCC | ..G...T..C......T... | NA | chr5:66830623-66830645 |
| PD-1_Ex_SD_OT10 | PD-1_Ex.1_SD_OT10 Rev 2 | GTCTCGTGGGCTCGGAGATGTGTATAAGAGACAGAGCCTTTCAGATTAGTCAGG | CAGCTATCTCAGAACCTTCC | ..G...T..C......T... | NA | chr5:66830623-66830645 |
| TRAC2_Ex.3_SA_OnT | TRAC2_Ex.3_SA_OnT Fwd 1 | TCGTCGGCAGCGTCAGATGTGTATAAGAGACAGTCTCAGAGCTTAGGATGCAC | TTCGTATcTGTAAAACCAAG | .................... | NA | chr14:23019485-23019507 |
| TRAC2_Ex.3_SA_OnT | TRAC2_Ex.3_SA_OnT Rev 1 | GTCTCGTGGGCTCGGAGATGTGTATAAGAGACAGTCTTGAAACACAATACTGTTGG | TTCGTATcTGTAAAACCAAG | .................... | NA | chr14:23019485-23019507 |
| TRAC2_Ex.3_SA_OnT | TRAC2_Ex.3_SA_OnT Fwd 2 | TCGTCGGCAGCGTCAGATGTGTATAAGAGACAGTCTCAGAGCTTAGGATGCAC | TTCGTATcTGTAAAACCAAG | .................... | NA | chr14:23019485-23019507 |
| TRAC2_Ex.3_SA_OnT | TRAC2_Ex.3_SA_OnT Rev 2 | GTCTCGTGGGCTCGGAGATGTGTATAAGAGACAGCTTGAAACACAATACTGTTGG | TTCGTATcTGTAAAACCAAG | .................... | NA | chr14:23019485-23019507 |
| TRAC2_Ex.3_SA_OT1 | TRAC2_Ex.3_SA_OT1 Fwd 1 | TCGTCGGCAGCGTCAGATGTGTATAAGAGACAGACATACATTGCCTTACTTTGC | TTGGGATCTTTAAAACCAAG | ..G.G....T.......... | NA | chr2:15232313-15232335 |
| TRAC2_Ex.3_SA_OT1 | TRAC2_Ex.3_SA_OT1 Rev 1 | GTCTCGTGGGCTCGGAGATGTGTATAAGAGACAGTTTTTGACTGCCAGAAGGT | TTGGGATCTTTAAAACCAAG | ..G.G....T.......... | NA | chr2:15232313-15232335 |
| TRAC2_Ex.3_SA_OT1 | TRAC2_Ex.3_SA_OT1 Fwd 2 | TCGTCGGCAGCGTCAGATGTGTATAAGAGACAGACATACATTGCCTTACTTTGC | TTGGGATCTTTAAAACCAAG | ..G.G....T.......... | NA | chr2:15232313-15232335 |
| TRAC2_Ex.3_SA_OT1 | TRAC2_Ex.3_SA_OT1 Rev 2 | GTCTCGTGGGCTCGGAGATGTGTATAAGAGACAGGGAAGCCAAAAGTTATACATGA | TTGGGATCTTTAAAACCAAG | ..G.G....T.......... | NA | chr2:15232313-15232335 |
| TRAC2_Ex.3_SA_OT2 | TRAC2_Ex.3_SA_OT2 Fwd 1 | TCGTCGGCAGCGTCAGATGTGTATAAGAGACAGAGTTTGGCATCTTCTTTACCT | TGAGCATCTGTAAAACCAAG | .GA.C............... | SYNE2 | chr14:64658071-64658093 |
| TRAC2_Ex.3_SA_OT2 | TRAC2_Ex.3_SA_OT2 Rev 1 | GTCTCGTGGGCTCGGAGATGTGTATAAGAGACAGAGTGGGGCTTCTCATCAC | TGAGCATCTGTAAAACCAAG | .GA.C............... | SYNE2 | chr14:64658071-64658093 |
| TRAC2_Ex.3_SA_OT2 | TRAC2_Ex.3_SA_OT2 Fwd 2 | TCGTCGGCAGCGTCAGATGTGTATAAGAGACAGAAGTTTGGCATCTTCTTTACC | TGAGCATCTGTAAAACCAAG | .GA.C............... | SYNE2 | chr14:64658071-64658093 |
| TRAC2_Ex.3_SA_OT2 | TRAC2_Ex.3_SA_OT2 Rev 2 | GTCTCGTGGGCTCGGAGATGTGTATAAGAGACAGAGTGGGGCTTCTCATCAC | TGAGCATCTGTAAAACCAAG | .GA.C............... | SYNE2 | chr14:64658071-64658093 |
| TRAC2_Ex.3_SA_OT3 | TRAC2_Ex.3_SA_OT3 Fwd 1 | TCGTCGGCAGCGTCAGATGTGTATAAGAGACAGAATGATAGATCCCAGCTGAA | TTCTAATCTCTAAAACCAAG | ...TA....C.......... | NA | chr11:116099208-116099230 |
| TRAC2_Ex.3_SA_OT3 | TRAC2_Ex.3_SA_OT3 Rev 1 | GTCTCGTGGGCTCGGAGATGTGTATAAGAGACAGCTTTCCTCCTTGCATGTATT | TTCTAATCTCTAAAACCAAG | ...TA....C.......... | NA | chr11:116099208-116099230 |
| TRAC2_Ex.3_SA_OT3 | TRAC2_Ex.3_SA_OT3 Fwd 2 | TCGTCGGCAGCGTCAGATGTGTATAAGAGACAGGGAGCCACAGATTAAATGAT | TTCTAATCTCTAAAACCAAG | ...TA....C.......... | NA | chr11:116099208-116099230 |
| TRAC2_Ex.3_SA_OT3 | TRAC2_Ex.3_SA_OT3 Rev 2 | GTCTCGTGGGCTCGGAGATGTGTATAAGAGACAGTCTTTCCTCCTTGCATGTAT | TTCTAATCTCTAAAACCAAG | ...TA....C.......... | NA | chr11:116099208-116099230 |
| TRAC2_Ex.3_SA_OT4 | TRAC2_Ex.3_SA_OT4 Fwd 1 | TCGTCGGCAGCGTCAGATGTGTATAAGAGACAGTCATGAATGGTGACTCAGAA | GTGGTATCTGCAAAACCAAG | G.G.......C......... | NA | chr3:83858142-83858164 |
| TRAC2_Ex.3_SA_OT4 | TRAC2_Ex.3_SA_OT4 Rev 1 | GTCTCGTGGGCTCGGAGATGTGTATAAGAGACAGAAATGCCAGCCACTTTTT | GTGGTATCTGCAAAACCAAG | G.G.......C......... | NA | chr3:83858142-83858164 |
| TRAC2_Ex.3_SA_OT4 | TRAC2_Ex.3_SA_OT4 Fwd 2 | TCGTCGGCAGCGTCAGATGTGTATAAGAGACAGAAATCATGAATGGTGACTCAG | GTGGTATCTGCAAAACCAAG | G.G.......C......... | NA | chr3:83858142-83858164 |
| TRAC2_Ex.3_SA_OT4 | TRAC2_Ex.3_SA_OT4 Rev 2 | GTCTCGTGGGCTCGGAGATGTGTATAAGAGACAGAAATGCCAGCCACTTTTT | GTGGTATCTGCAAAACCAAG | G.G.......C......... | NA | chr3:83858142-83858164 |
| TRAC2_Ex.3_SA_OT5 | TRAC2_Ex.3_SA_OT5 Fwd 1 | TCGTCGGCAGCGTCAGATGTGTATAAGAGACAGGCCAAACCATATTAGCAAAC | TCCTCATGTGTAAAACCAAG | .C.TC..G............ | NA | chr10:33362901-33362923 |
| TRAC2_Ex.3_SA_OT5 | TRAC2_Ex.3_SA_OT5 Rev 1 | GTCTCGTGGGCTCGGAGATGTGTATAAGAGACAGTTGAGTTCATGAGAATCGTG | TCCTCATGTGTAAAACCAAG | .C.TC..G............ | NA | chr10:33362901-33362923 |
| TRAC2_Ex.3_SA_OT5 | TRAC2_Ex.3_SA_OT5 Fwd 2 | TCGTCGGCAGCGTCAGATGTGTATAAGAGACAGGCCAAACCATATTAGCAAAC | TCCTCATGTGTAAAACCAAG | .C.TC..G............ | NA | chr10:33362901-33362923 |
| TRAC2_Ex.3_SA_OT5 | TRAC2_Ex.3_SA_OT5 Rev 2 | GTCTCGTGGGCTCGGAGATGTGTATAAGAGACAGATTGAGTTCATGAGAATCGTG | TCCTCATGTGTAAAACCAAG | .C.TC..G............ | NA | chr10:33362901-33362923 |
| TRAC2_Ex.3_SA_OT6 | TRAC2_Ex.3_SA_OT6 Fwd 1 | TCGTCGGCAGCGTCAGATGTGTATAAGAGACAGGCAAACGTACACTGTAATGC | TCTGCATCTTTAAAACCAAG | .CT.C....T.......... | GRIA1 | chr5:153014461-153014483 |
| TRAC2_Ex.3_SA_OT6 | TRAC2_Ex.3_SA_OT6 Rev 1 | GTCTCGTGGGCTCGGAGATGTGTATAAGAGACAGGCTTTGCTGAGACCATAGAT | TCTGCATCTTTAAAACCAAG | .CT.C....T.......... | GRIA1 | chr5:153014461-153014483 |
| TRAC2_Ex.3_SA_OT6 | TRAC2_Ex.3_SA_OT6 Fwd 2 | TCGTCGGCAGCGTCAGATGTGTATAAGAGACAGGCAAACGTACACTGTAATGC | TCTGCATCTTTAAAACCAAG | .CT.C....T.......... | GRIA1 | chr5:153014461-153014483 |
| TRAC2_Ex.3_SA_OT6 | TRAC2_Ex.3_SA_OT6 Rev 2 | GTCTCGTGGGCTCGGAGATGTGTATAAGAGACAGATGCTTTGCTGAGACCATAG | TCTGCATCTTTAAAACCAAG | .CT.C....T.......... | GRIA1 | chr5:153014461-153014483 |
| TRAC2_Ex.3_SA_OT7 | TRAC2_Ex.3_SA_OT7 Fwd 1 | TCGTCGGCAGCGTCAGATGTGTATAAGAGACAGCAAAGTGCTGGGATTACAGA | TTTGTATCTTTAAAACCATG | ..T......T........T. | TOPBP1 | chr3:133341480-133341502 |
| TRAC2_Ex.3_SA_OT7 | TRAC2_Ex.3_SA_OT7 Rev 1 | GTCTCGTGGGCTCGGAGATGTGTATAAGAGACAGTGCAAAGTTTTATGTAGTTTAAGTG | TTTGTATCTTTAAAACCATG | ..T......T........T. | TOPBP1 | chr3:133341480-133341502 |
| TRAC2_Ex.3_SA_OT7 | TRAC2_Ex.3_SA_OT7 Fwd 2 | TCGTCGGCAGCGTCAGATGTGTATAAGAGACAGCAAAGTGCTGGGATTACAGA | TTTGTATCTTTAAAACCATG | ..T......T........T. | TOPBP1 | chr3:133341480-133341502 |
| TRAC2_Ex.3_SA_OT7 | TRAC2_Ex.3_SA_OT7 Rev 2 | GTCTCGTGGGCTCGGAGATGTGTATAAGAGACAGAATTTGCAAAGTTTTATGTAGTTT | TTTGTATCTTTAAAACCATG | ..T......T........T. | TOPBP1 | chr3:133341480-133341502 |
| TRAC2_Ex.3_SA_OT8 | TRAC2_Ex.3_SA_OT8 Fwd 1 | TCGTCGGCAGCGTCAGATGTGTATAAGAGACAGTGGGACTCTTGGTTCTGTAT | TTCTTATGTGTGAAACCAAG | ...T...G...G........ | NTNG1 | chr1:107907252-107907274 |
| TRAC2_Ex.3_SA_OT8 | TRAC2_Ex.3_SA_OT8 Rev 1 | GTCTCGTGGGCTCGGAGATGTGTATAAGAGACAGTTTTTGTTGTTGTTTTACTTGAA | TTCTTATGTGTGAAACCAAG | ...T...G...G........ | NTNG1 | chr1:107907252-107907274 |
| TRAC2_Ex.3_SA_OT8 | TRAC2_Ex.3_SA_OT8 Fwd 2 | TCGTCGGCAGCGTCAGATGTGTATAAGAGACAGTTGGGACTCTTGGTTCTGTA | TTCTTATGTGTGAAACCAAG | ...T...G...G........ | NTNG1 | chr1:107907252-107907274 |
| TRAC2_Ex.3_SA_OT8 | TRAC2_Ex.3_SA_OT8 Rev 2 | GTCTCGTGGGCTCGGAGATGTGTATAAGAGACAGTTTTTGTTGTTGTTTTACTTGAA | TTCTTATGTGTGAAACCAAG | ...T...G...G........ | NTNG1 | chr1:107907252-107907274 |
| TRAC2_Ex.3_SA_OT9 | TRAC2_Ex.3_SA_OT9 Fwd 1 | TCGTCGGCAGCGTCAGATGTGTATAAGAGACAGAAATTCTTCTTGGGCTCAG | CTCTCTTCTGTAAAACCAAG | C..TCT.............. | KCNQ5 | chr6:73675659-73675681 |
| TRAC2_Ex.3_SA_OT9 | TRAC2_Ex.3_SA_OT9 Rev 1 | GTCTCGTGGGCTCGGAGATGTGTATAAGAGACAGGCTCATTACTGGGTTAAGCA | CTCTCTTCTGTAAAACCAAG | C..TCT.............. | KCNQ5 | chr6:73675659-73675681 |
| TRAC2_Ex.3_SA_OT9 | TRAC2_Ex.3_SA_OT9 Fwd 2 | TCGTCGGCAGCGTCAGATGTGTATAAGAGACAGGAAATTCTTCTTGGGCTCAG | CTCTCTTCTGTAAAACCAAG | C..TCT.............. | KCNQ5 | chr6:73675659-73675681 |
| TRAC2_Ex.3_SA_OT9 | TRAC2_Ex.3_SA_OT9 Rev 2 | GTCTCGTGGGCTCGGAGATGTGTATAAGAGACAGGCTCATTACTGGGTTAAGCA | CTCTCTTCTGTAAAACCAAG | C..TCT.............. | KCNQ5 | chr6:73675659-73675681 |
| TRAC2_Ex.3_SA_OT10 | TRAC2_Ex.3_SA_OT10 Fwd 1 | TCGTCGGCAGCGTCAGATGTGTATAAGAGACAGTCTAGGCTCTTGACACCATC | TGGGTTTCTTTAAAACCAAG | .GG..T...T.......... | FSHR | chr2:49345982-49346004 |
| TRAC2_Ex.3_SA_OT10 | TRAC2_Ex.3_SA_OT10 Rev 1 | GTCTCGTGGGCTCGGAGATGTGTATAAGAGACAGGAACAACTAGACCCAATGTGA | TGGGTTTCTTTAAAACCAAG | .GG..T...T.......... | FSHR | chr2:49345982-49346004 |
| TRAC2_Ex.3_SA_OT10 | TRAC2_Ex.3_SA_OT10 Fwd 2 | TCGTCGGCAGCGTCAGATGTGTATAAGAGACAGGTCTAGGCTCTTGACACCAT | TGGGTTTCTTTAAAACCAAG | .GG..T...T.......... | FSHR | chr2:49345982-49346004 |
| TRAC2_Ex.3_SA_OT10 | TRAC2_Ex.3_SA_OT10 Rev 2 | GTCTCGTGGGCTCGGAGATGTGTATAAGAGACAGGAACAACTAGACCCAATGTGA | TGGGTTTCTTTAAAACCAAG | .GG..T...T.......... | FSHR | chr2:49345982-49346004 |
