## Supplementary material for "Highly efficient multiplex human T cell engineering without double-strand breaks using Cas9 base editors"

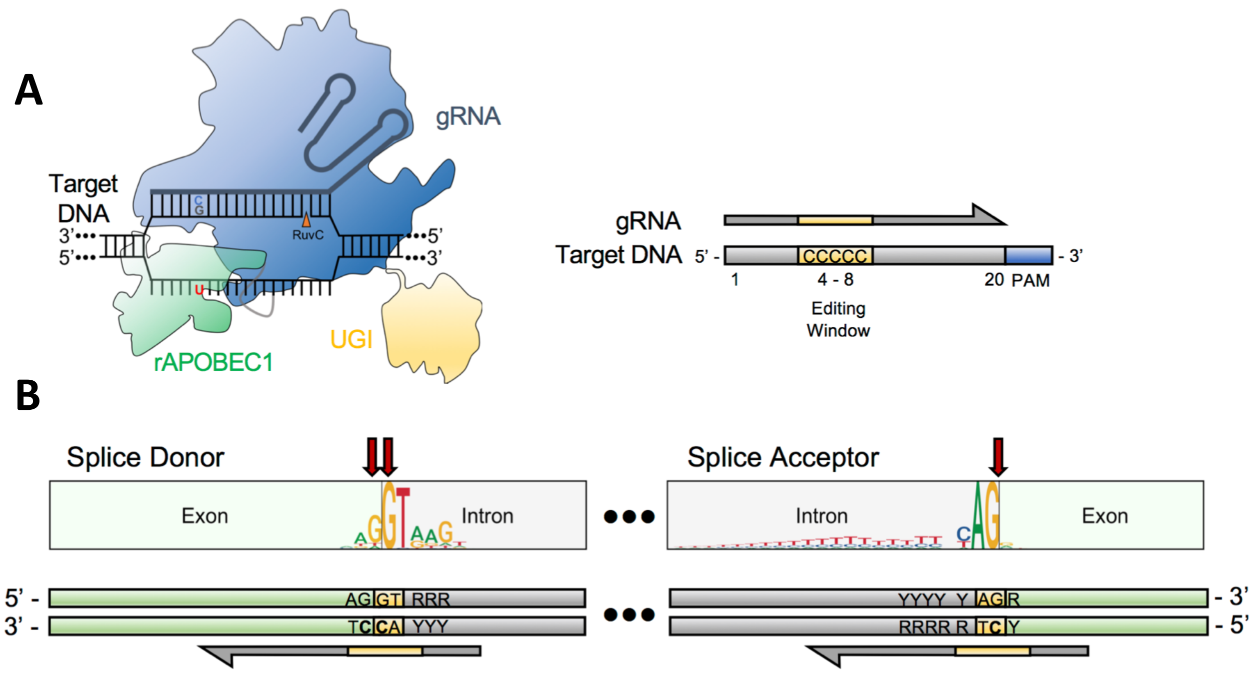


**Supplemental Data 1**: **Targeting splice sites using base editor. A.** Diagram depicting Cas9 base editor (BE) bound to target DNA (left) and protospacer depicting the purported base editing window achieved with BE3 and BE4 (right). **B.** Logo diagrams depicting the consensus sequence of mammalian splice donor (SD) and splice acceptor (SA) elements and the related orientation of protospacers utilized for BE knockout via splice site disruption.

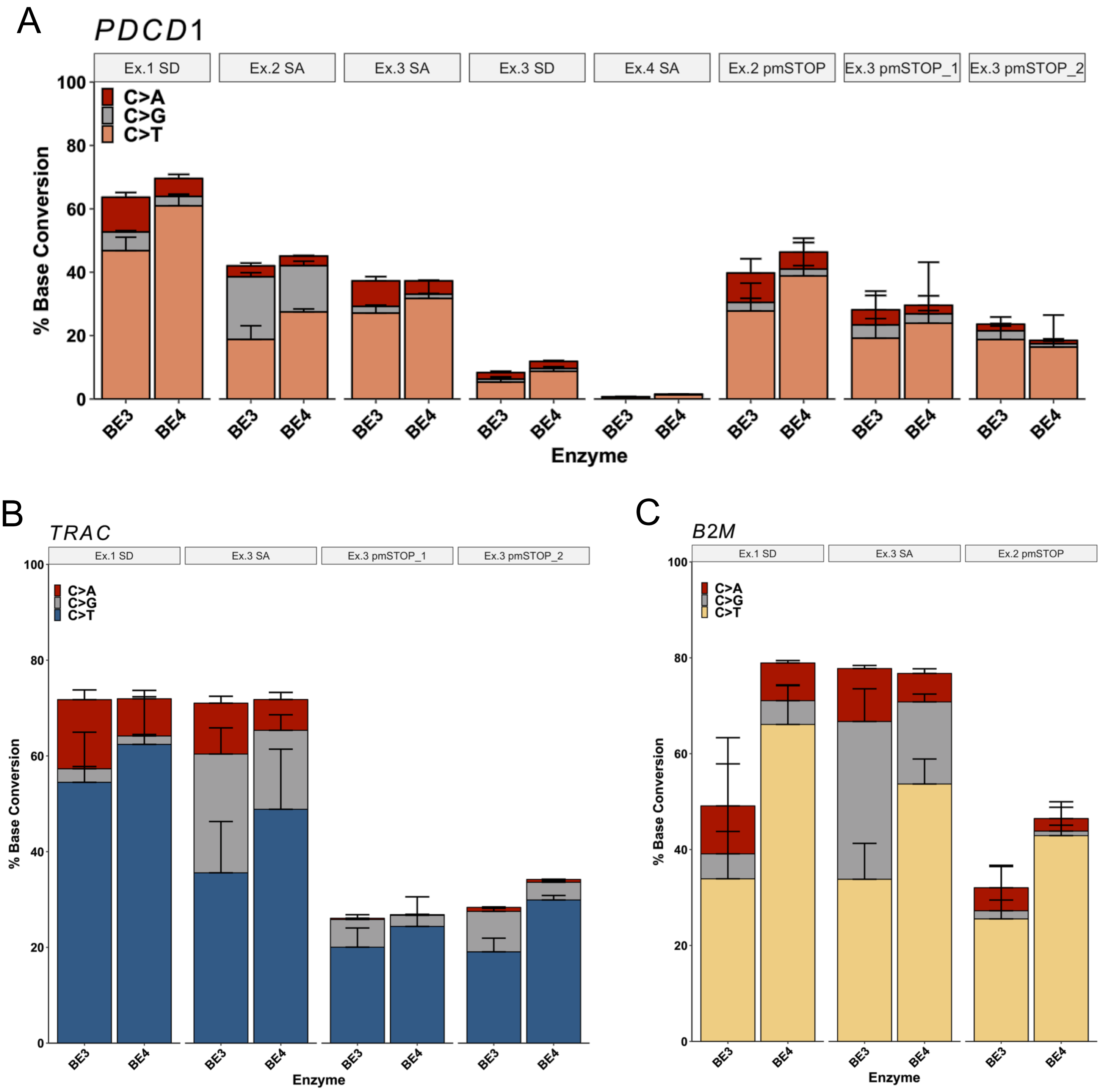


**Supplemental Data 2. Non-target editing for each sgRNA in figure 1.** Data is analyzed from NGS. Height of stacked bars represents mean, with error bars ± 1 standard deviation. *n=3 independent donors.*

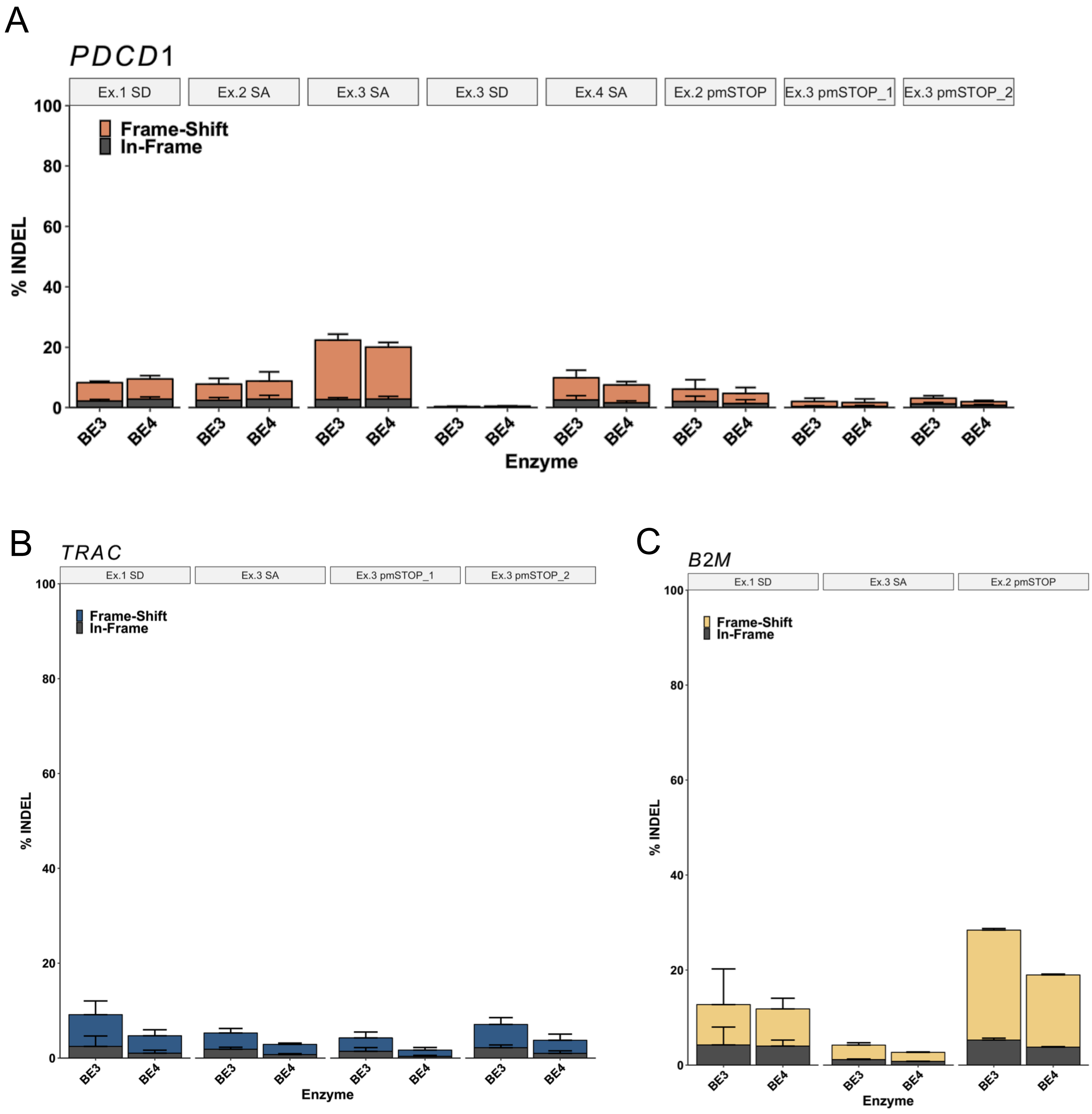


**Supplemental Data 3. Indels for all samples in Figure 1.** Data is analyzed from NGS. Height of stacked bars represents mean, with error bars ± 1 standard deviation. *n=3 independent donors.*

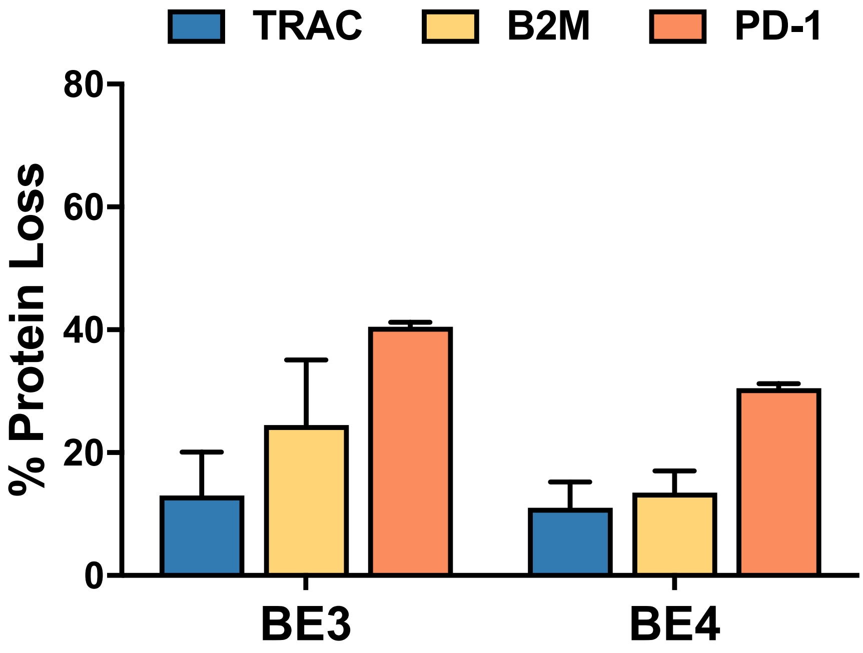


**Supplemental Data 4. Multiplex base editing of T cells using first-generation, low-dose (1.5 µg) BE3 or BE4 mRNA.** Bar graph depicting base editor mediated knockout of TRAC, B2M and PD-1 at the protein level. Protein expression was assessed via flow cytometry as described in the methods section. *n = 2 independent donors.*

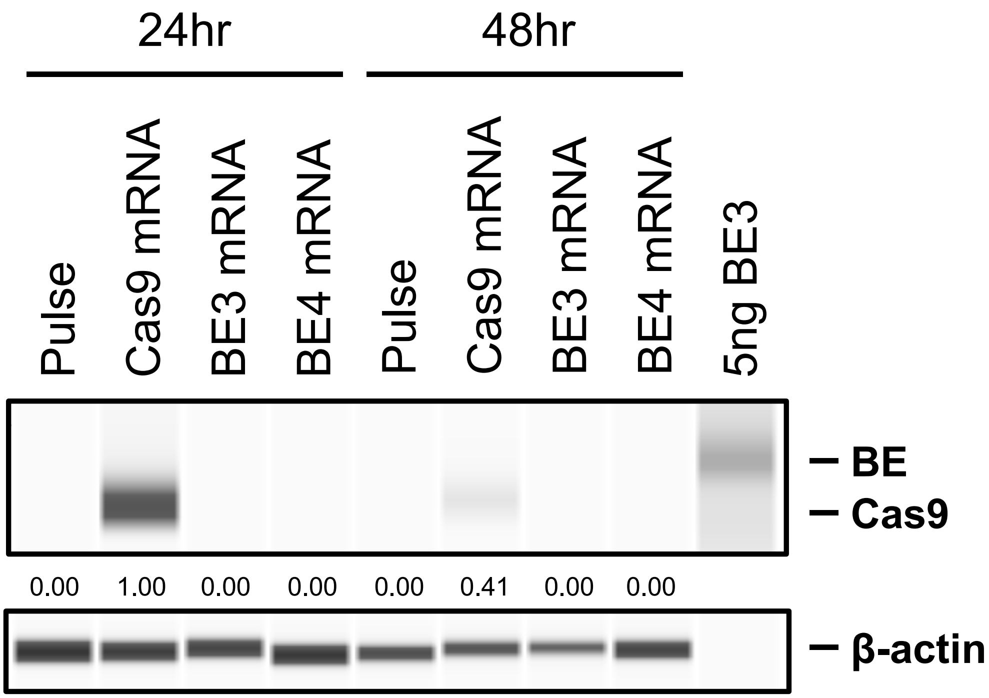


**Supplemental Data 5. Base editor protein levels following electroporation of T cells.** Digital western blot results assessing the protein level achieved using mRNA encoding Cas9, BE3, and BE4 at 24hrs and 48hrs post electroporation of stimulated T cells. Purified BE3 protein was also used as a positive control for antibody detection of BE protein.

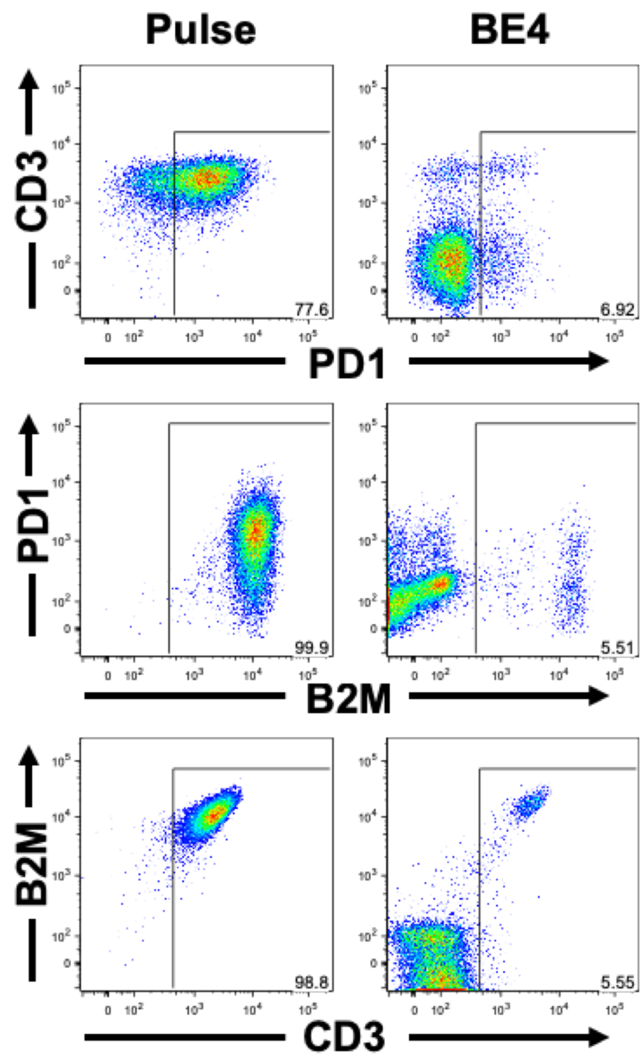


**Supplemental Data 6. Representative flow plots of PD1, B2M and TRAC upon re-stimulation.** Five days post electroporation T cells are re-stimulated to induce expression of PD-1, allowing for the assessment of PD-1 protein knockout frequencies. Shown here are representative flow cytometry plots of TRAC, B2M and PD-1 expression of donor-matched T cell following multiplex coBE4 mRNA editing and pulse-only control (*left column*).

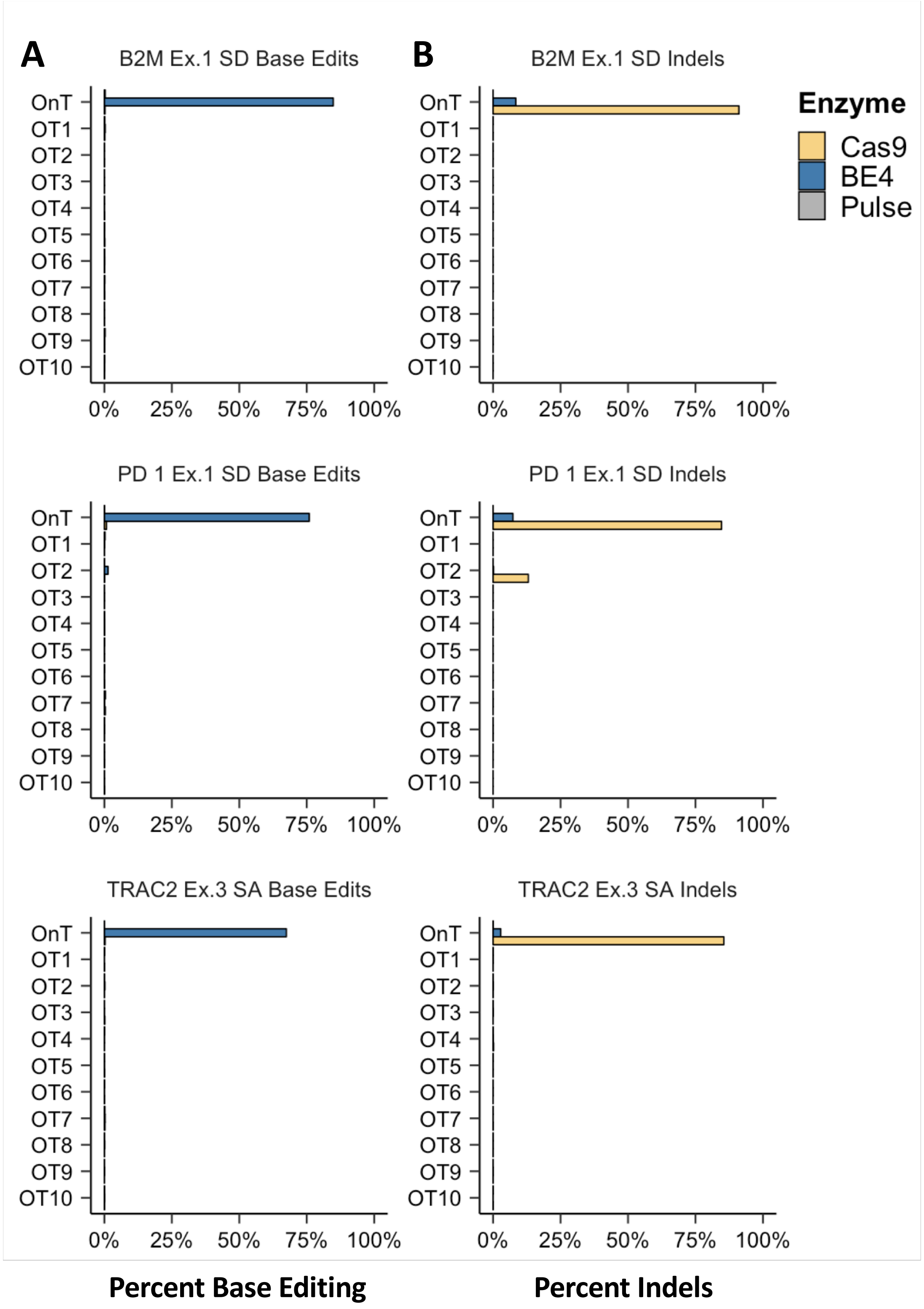


**Supplemental Data 7: Assessment of computationally predicted off-target base editing and indel formation.** Base editing **(A)** and indel **(B)** frequency at on-target and top 10 computationally predicted off-target sgRNA binding sites, assessed using next generation sequencing, using optimal sgRNAs targeting *TRAC*, *B2M* or *PDCD1* combined with Cas9 or BE4 mRNA in T cells.

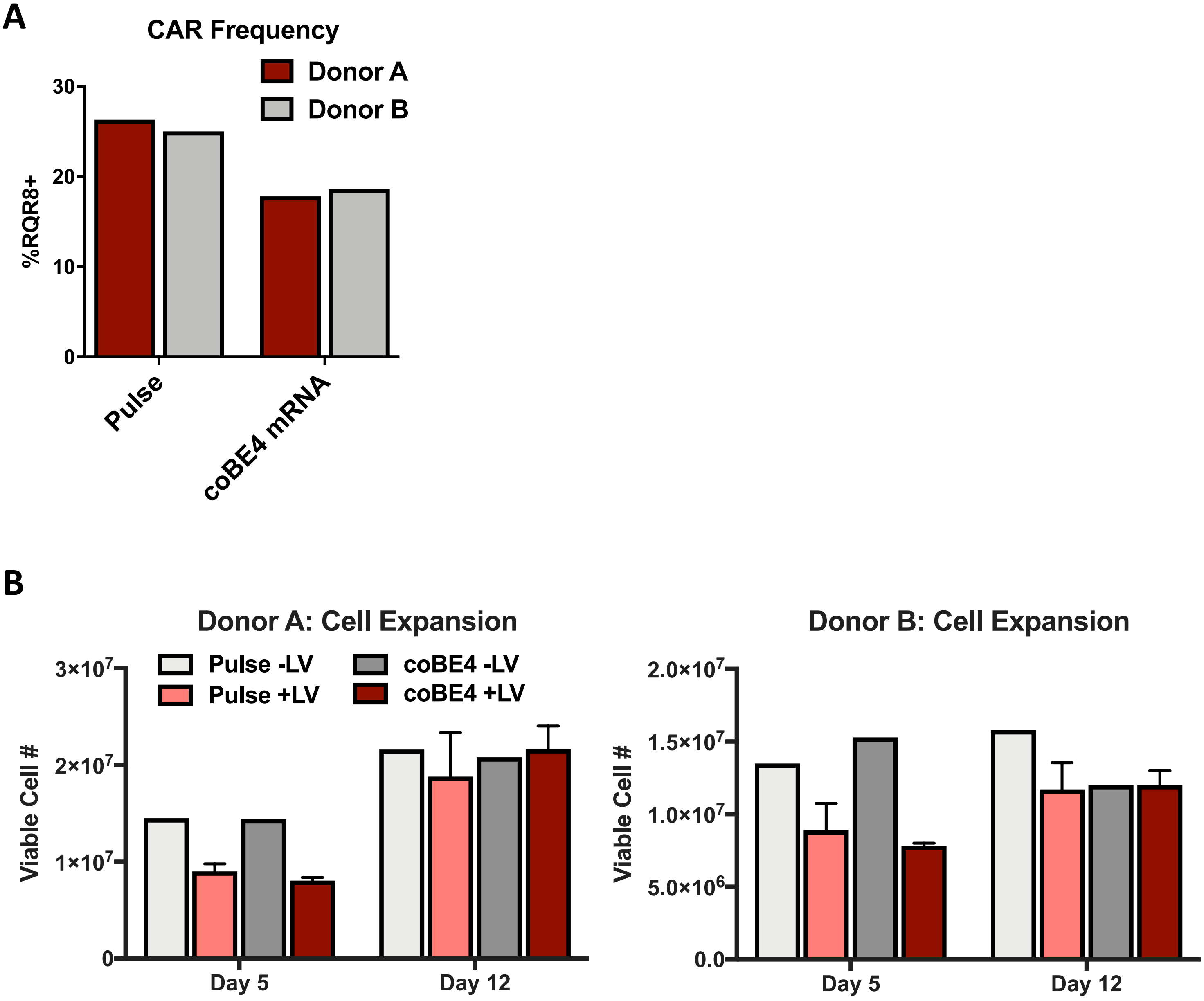


**Supplemental Data 8: CAR transduction and T cell expansion efficiency. A.** Bar graph depicting the frequency of transduced T cells using the MND-CD19 CAR-RQR8 lentiviral vectors, MOI of 20, via staining for RQR8 in two independent donors. RQR8 is a hybrid molecule containing domains for staining with CD34 and CD20 specific antibody and serves as a surrogate for determining CAR positive T cell frequency. **B.** Bar graphs depicting the number of viable cells at day 5 and 12 post electroporation and transduction. *n=2 independent donors.*

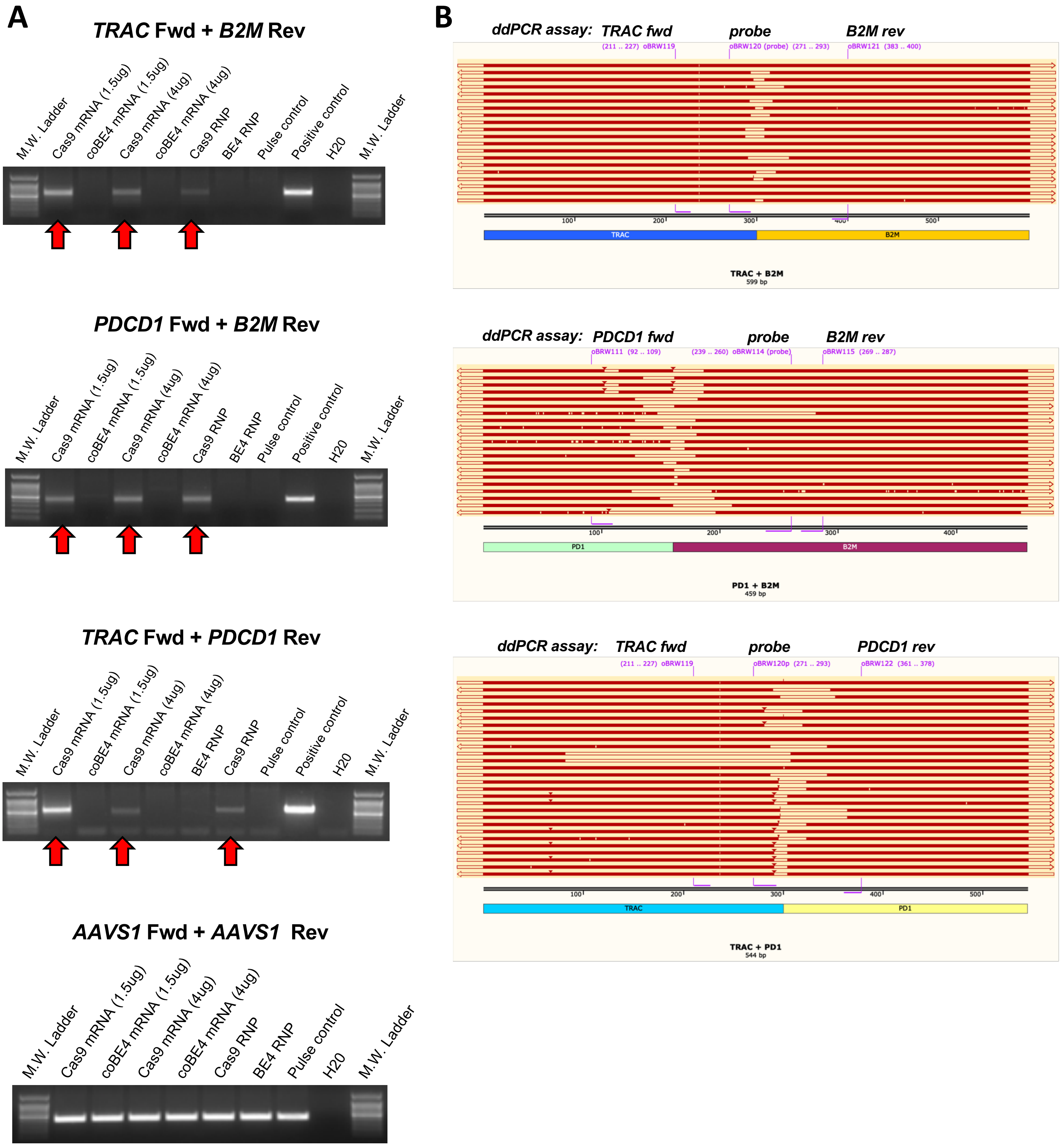


**Supplemental Data 9. Sequencing of sub-cloned PCR products spanning translocation junctions. A.** Results of translocation PCR performed between the noted target genes using Cas9 or BE mRNA or protein, as noted. An AAVS1 control PCR was also performed to confirm gDNA quality and functionality for PCR**. B.** PCR products from (A) were TA cloned into TOPO plasmids and subsequently analyzed via Sanger sequencing. Resultant chromatograms were then aligned to a hypothetical 'perfect' junction sequence between the noted target gene gRNA cut sites and aligned. Also depicted are ddPCR probes used to generate the data presented in main figure 3.
